# Metabolic Reprogramming of the Neovascular Niche Promotes Regenerative Angiogenesis in Proliferative Retinopathy

**DOI:** 10.1101/2023.11.13.566898

**Authors:** Gael Cagnone, Sheetal Pundir, Jin Sung-Kim, José Carlos Rivera, Tapan Agnihotri, Noémie-Rose Harvey, Emilie Heckel, Charlotte Betus, Mei Xi Chen, Anu Situ, Perrine Gaub, Nicholas Kim, Ashim Das, Severine Leclerc, Florian Wünnemann, Gregor Andelfinger, Sergio Crespo-Garcia, Alexandre Dubrac, Flavio Rezende, Clary B. Clish, Bruno Maranda, Lois E. H. Smith, Przemyslaw Sapieha, Jean-Sébastien Joyal

## Abstract

Healthy blood vessels supply neurons to preserve metabolic function. In blinding ischemic proliferative retinopathies (PRs), pathological neovascular tufts often emerge in lieu of needed physiological neuroretina revascularization. We show that metabolic shifts in the neurovascular niche define this angiogenic dichotomy between healthy and diseased blood vessel growth. Fatty acid oxidation (FAO) metabolites accumulated in human and murine retinopathy samples. Neovascular tufts with a distinct single-cell transcriptional signature highly expressed FAO enzymes. The deletion of *Sirt3*, an FAO regulator, shifted the neurovascular niche metabolism from FAO to glycolysis and suppressed tuft formation. This metabolic transition increased *Vegf* expression in astrocytes and reprogrammed pathological EC to a physiological phenotype, hastening vascular regeneration of the ischemic retina. Our findings identify SIRT3 as a metabolic switch in the neurovascular niche, offering a new therapeutic target for optimizing ischemic tissue revascularization.

**Highlights:**

1. Pathological EC favor FAO over glycolysis.
2. Unique signature for pathological EC found in proliferative retinopathy model.
3. *Sirt3* deletion shifts astrocytes and EC metabolism from FAO to glycolysis.
4. Metabolic reprogramming of the vascular niche enhances physiological revascularization.

**Graphical Abstract:** 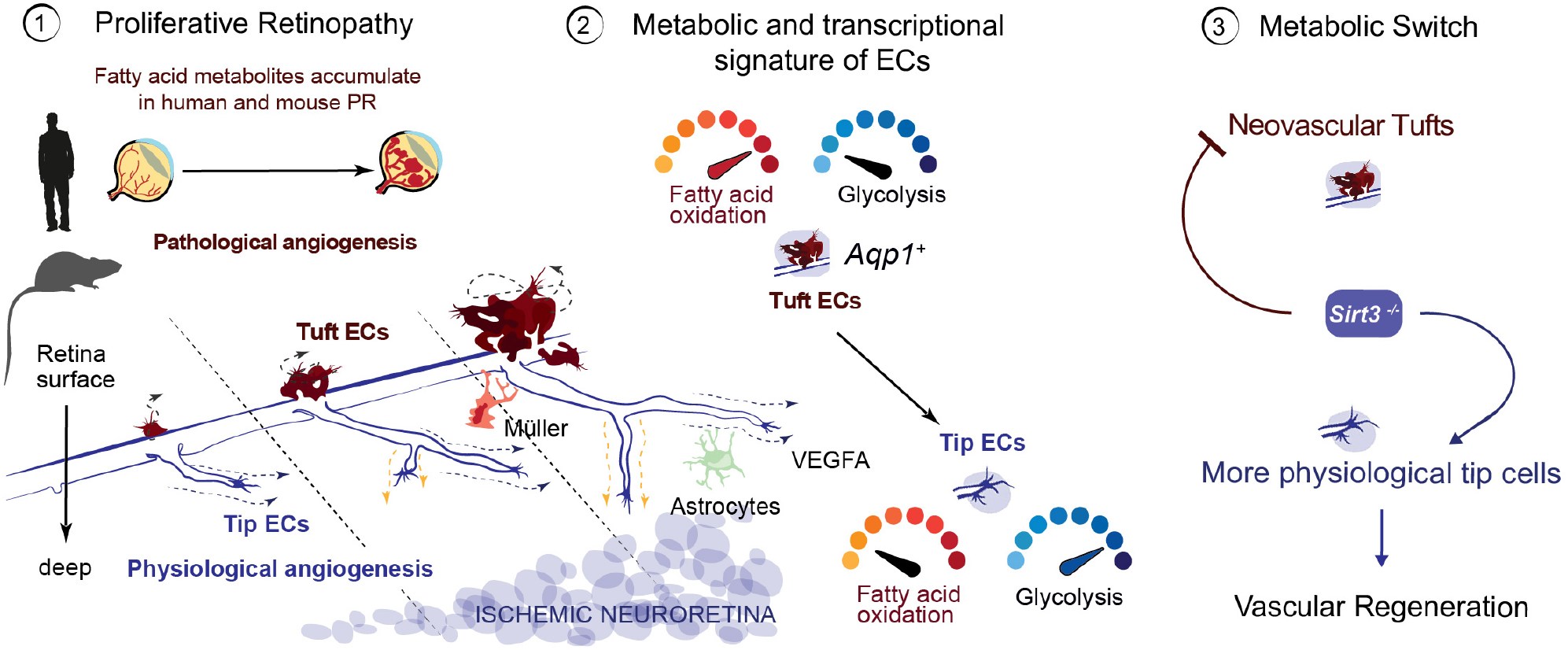

## INTRODUCTION

Proliferative retinopathy (PR) is the leading cause of blindness in children with retinopathy of prematurity and working-age adults with diabetic retinopathy^1^. The initial blood vessel loss in PR and the neuroretinal ischemia that follows trigger the excessive and misplaced production of vascular endothelial growth factor (VEGF), driving pathological angiogenesis in an attempt to reinstate metabolic homeostasis^1,2^. While inhibiting VEGF to block pathological neovascular tuft formation has been the focus of therapeutic interventions to date, the regenerative forces driving physiological revascularization of the ischemic neuroretina have been less explored. Discerning unique molecular fingerprints for pathological tufts and physiological angiogenesis, along with their associated vascular niche, could help reveal the drivers of healthy revascularization and neuronal survival.

Neuroglial cells of the vascular unit secrete growth factors and guidance cues that shape vascular architecture to meet their metabolic requirements^3,4^. During vascular development, astrocytes^5,6^ and retinal ganglion cells^7^ (RGCs) secrete VEGF that guides the advancing migratory tip EC of the vascular front. During PR, when the retina has few blood vessels and is hypoxic and nutrient starved, excessive VEGF and repulsive guidance cues are secreted by ischemic neuroglial cells^7–10^. However, it is unclear whether metabolic changes in the vascular niche drive angiogenic fates, be they healthy or diseased.

During developmental angiogenesis, EC rely primarily on glycolysis to produce energy^11^. Glycolysis contributes to EC proliferation and migration required for vessel growth. Glycolytic enzymes are localized in lamellipodia and leading membrane ruffles of EC, powering the migration of endothelial tip cells^12^ that guide physiological vascular growth. Since glucose diffuses further from vessels than oxygen^12–14^, EC can rely on anaerobic glycolysis when sprouting in avascular tissues. Pharmacological inhibition of glycolysis prevents sprouting angiogenesis^12,15^. Hence, glycolysis may be a metabolic trait of EC that would be advantageous to revascularizing the ischemic retina.

Fatty acid oxidation (FAO) also promotes EC proliferation^11^. In addition to energy production, fatty acids are an essential carbon source to produce *de novo* nucleotides for DNA replication in EC^16^. This biosynthetic function contributes to EC proliferation, but unlike glycolysis, it is not associated with increased EC migration, a characteristic of tip cells driving vascular regeneration^12^. Inhibition of carnitine palmitoyl-transferase 1 (CPT1), shuttling long chain fatty acids inside mitochondria for oxidation, decreases pathological tufts in murine PR^16^, but its impact on physiological revascularisation is unknown.

Sirtuins (SIRT) are NAD+-dependent deacetylases pivotal for mitochondrial function^17^. SIRT3 is a master regulator of FAO and oxidative energy metabolism within mitochondria^18–20^. Since both FAO and glycolysis fuel many critical biological functions, including angiogenesis, we reasoned that SIRT3 could be a metabolic switch able to modulate the relative balance of energy utilization within the vascular niche without the deleterious effects of completely abrogating one of these essential energy sources. We combined unbiased large-scale metabolomics and single-cell transcriptomics data to define distinct metabolic and transcriptional signatures for physiological and pathological EC. While glycolysis is essential to the migratory and regenerative tip EC, we show that FAO is a hallmark of misguided yet actively proliferating pathological neovascular EC. Metabolic reprogramming of the retinal microenvironment caused by *Sirt3* deletion shifts neovascular metabolism from FAO to glycolysis and enhances physiological vascular regeneration of the retina.

## RESULTS

### Fatty-acid oxidation is a metabolic hallmark of pathological angiogenesis

We first examined the metabolite profile of vitreous humor from subjects with proliferative diabetic retinopathy (Table S1, Figures S1A-C). Vitreous samples were collected adjacent to leaky pathological neovascular tufts that characterize PR (Figure 1A). Amongst the top 25 most significantly dysregulated metabolites, we observed an accumulation of acylcarnitines in human neovascular PR samples compared to control subjects with epiretinal membranes (Figure 1A, red font). Fatty acylcarnitines are by-products of fatty-acid β-oxidation (FAO), the process of oxidizing fatty acids in mitochondria to produce acetyl-CoA (Figure 1B). Using Metaboanalyst^21^, a metabolomics analysis tool, 5 of the 6 most enriched metabolic pathways in human PR pertained to FA metabolism, including long and short-chain saturated FAO pathways (Figure 1B). Hence, the accumulation of FAO metabolites is a prominent characteristic of human proliferative retinopathy.

**Figure 1.**
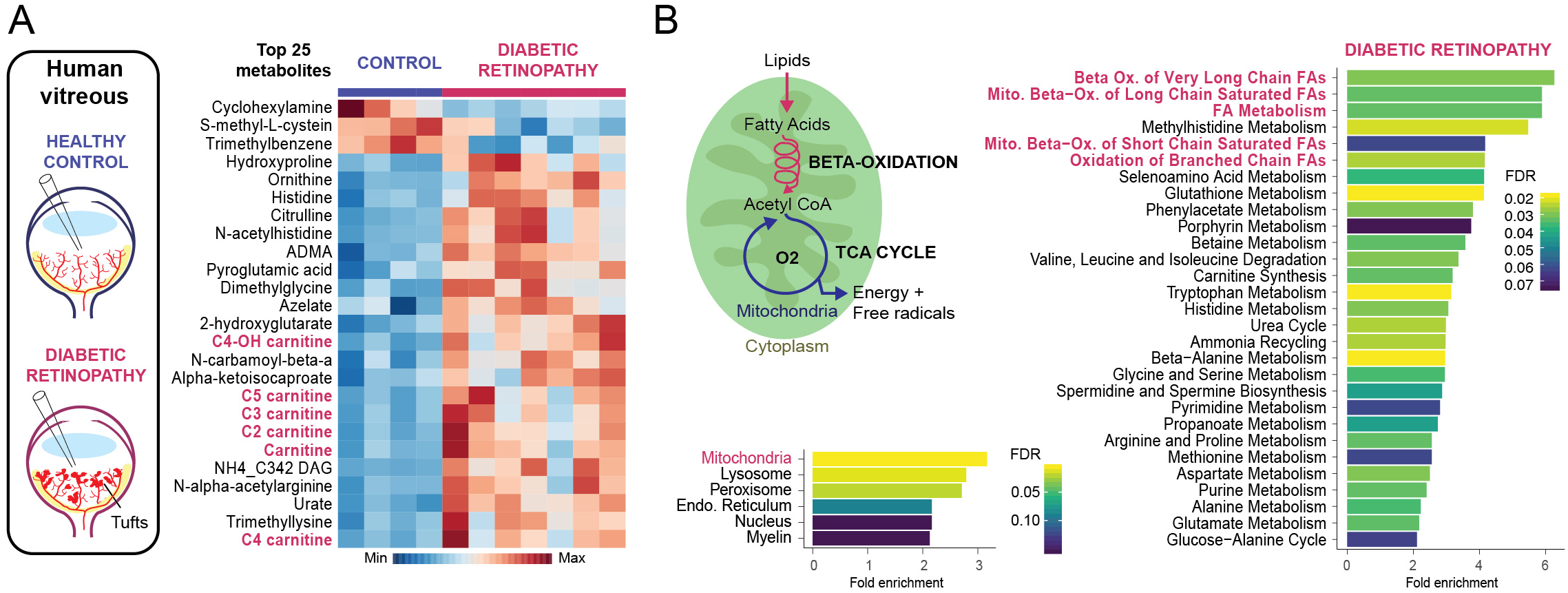
Fatty-acid oxidation is a hallmark of human proliferative diabetic retinopathy. **A.** Heatmap of the top 25 most dysregulated metabolites from vitreous biopsies of subjects with epiretinal membranes (Control, n=4) and proliferative diabetic retinopathy (n=7) by LC/MS/MS. **B.** Metabolite set enrichment analysis of dysregulated metabolites for metabolic pathways and intracellular localisations. Statistical significance was established using a two-tailed Student’s *t*-test, P value < 0.05 See also Figure S1 and Table S1.

Vitreous metabolites are often used as surrogate markers of adjacent retinal metabolism^22,23^. However, decerning the precise cellular origin of these metabolites in human retinas could impair residual vision. Fibrovascular membranes (FVM) resected from diabetic retinopathy patients have been analyzed through single-cell RNAseq^24^ revealing some tip-like endothelial cells (FVM EC 2, *ESM1*). Still, these lesions typically lack active neovascular tufts (Figure S1D-F). Consequently, we adopted the oxygen-induced retinopathy (OIR) model, which simulates the formation of pathological neovascular tufts characteristic of PR (Figure 2A)^25^. In this model, mice pups are exposed to high oxygen concentration for 5 days (post-natal day (P) 7 to 12), causing vaso-obliteration (VO) and subsequent retinal ischemia. After returning to room air (P12), the ischemic neuroretina triggers both its physiological revascularization and the formation of pathological neovascular tufts that invade the vitreous^25^. At the peak of tuft formation in mouse PR (P17), we observed comparable enrichments of FAO metabolites to human PR (Figures 2B and S2A-C). Thus, the metabolomic profile in mice with PR closely resembles the human condition, establishing a relevant model to investigate the influence of FAO on neovascular diseases.

**Figure 2.**
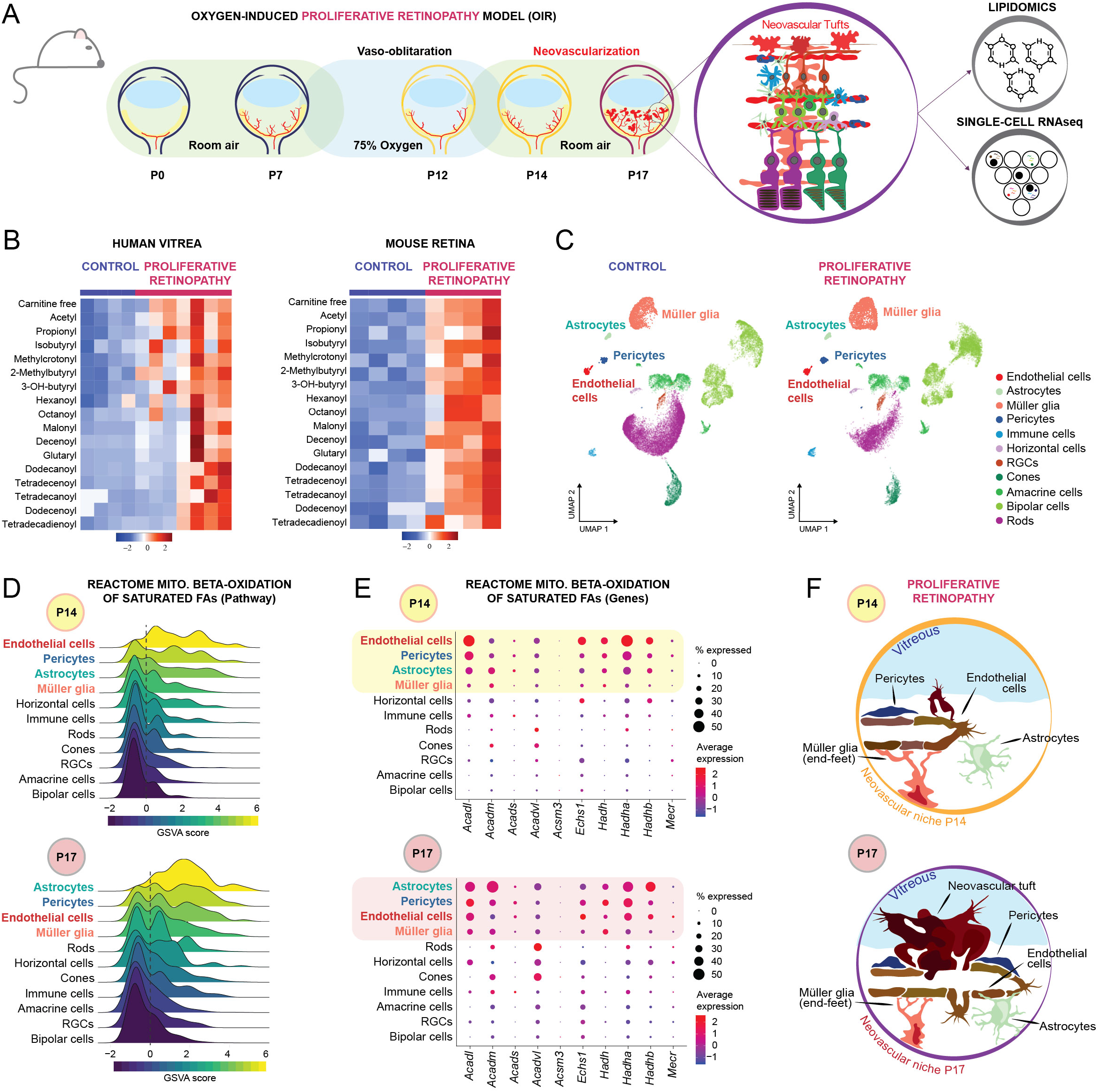
FAO is enriched in the neurovascular unit in mouse proliferative retinopathy. **A.** Schematic representation of the oxygen-induced retinopathy (OIR) mouse model of proliferative retinopathy. **B.** Heatmap of fatty acylcarnitine metabolites measured in human vitreous and mouse retinas of control (human n = 4, mouse n = 4) and proliferative retinopathy samples (human n = 7, mouse n = 4). **C.** UMAP of single-cell RNAseq from normoxic (n=6, 21305 cells) and OIR (n=8, 17814 cells) retinas taken during the neovascularisation period (P14-P17) representing the 11 retinal cell types identified by graph-based clustering of normalized RNA count (GEO accession number GSE150703). **D.** Ridge plot of GSVA score for REACTOME mitochondrial beta-oxidation of saturated fatty acids pathway for OIR cell types at P14 and P17. **E.** Dot plot illustrating the expression levels of genes from the REACTOME pathway related to mitochondrial beta-oxidation of saturated fatty acids across OIR cell types at P14 and P17. **F.** Graphical representation of the vascular unit during the neovascular phase of OIR. See also Figure S2.

To determine the retinal source of FAO intermediates in PR, we compared the transcriptome of individual retinal cells from mice exposed to OIR and their normoxic littermate controls at the instigation of retinal revascularization (at P14) and the peak of pathological angiogenesis (at P17). Retinal cells were annotated, as previously described^26^ (Figure 2C). Gene set variation analysis (GSVA) for FAO pathways showed that cell types forming the vascular unit (EC, pericytes, astrocytes, and Müller glia) were the most enriched in transcripts coding for saturated FAO enzymes at both time points in the OIR hypoxic retina (Figure 2D). EC (P14) and astrocytes (P17) showed strong transcriptional expression of mitochondrial FAO genes (*Acadl, Acadvl, Hadh, Hadha,* and *Hadhb*) compared to other cell types (Figures 2E and S2D). These results indicated that cell types comprising the neovascular unit (Figure 2F), particularly EC and astrocytes, may contribute to the accumulation of FAO intermediates in ischemic PR.

### Single-cell RNAseq identifies a unique transcriptional signature for neovascular tufts

Uncovering the specific molecular fingerprints of pathological neovessels has been a critical objective in vascular biology. To examine EC heterogeneity in PR, we enriched retinal single-cell suspensions for EC (using magnetic beads coated with PECAM1) and performed a single-cell transcriptomic analysis of OIR and normoxic retinas (Figures S3A-B). The unbiased assessment of uniform manifold approximation and projection (UMAP) results uncovered six retinal EC subclusters (Figure 3A) comprising five known EC subtypes according to vascular zonation (arterial, vein, capillary, proliferative, tip cells^27^) and one previously unidentified EC subtype (*Aqp1+* EC, Figure 3B). Normoxic control retinas were composed primarily of capillary, vein, and arterial EC, with relatively few tip cells (1-3%) and proliferating EC (0-5%). In contrast, OIR retinas at P14, following the initial vaso-obliterative phase, contained more proliferative (24%) and endothelial tip cells (25%) to revascularize the ischemic retina and less mature arteries (2%) and veins (32%) (Figure 3C). Later, at P17, once further retinal revascularization was achieved, arterial (11%) and vein EC (57%) were more abundant, and fewer tip (10%) and proliferative EC (5%) were recovered.

**Figure 3.**
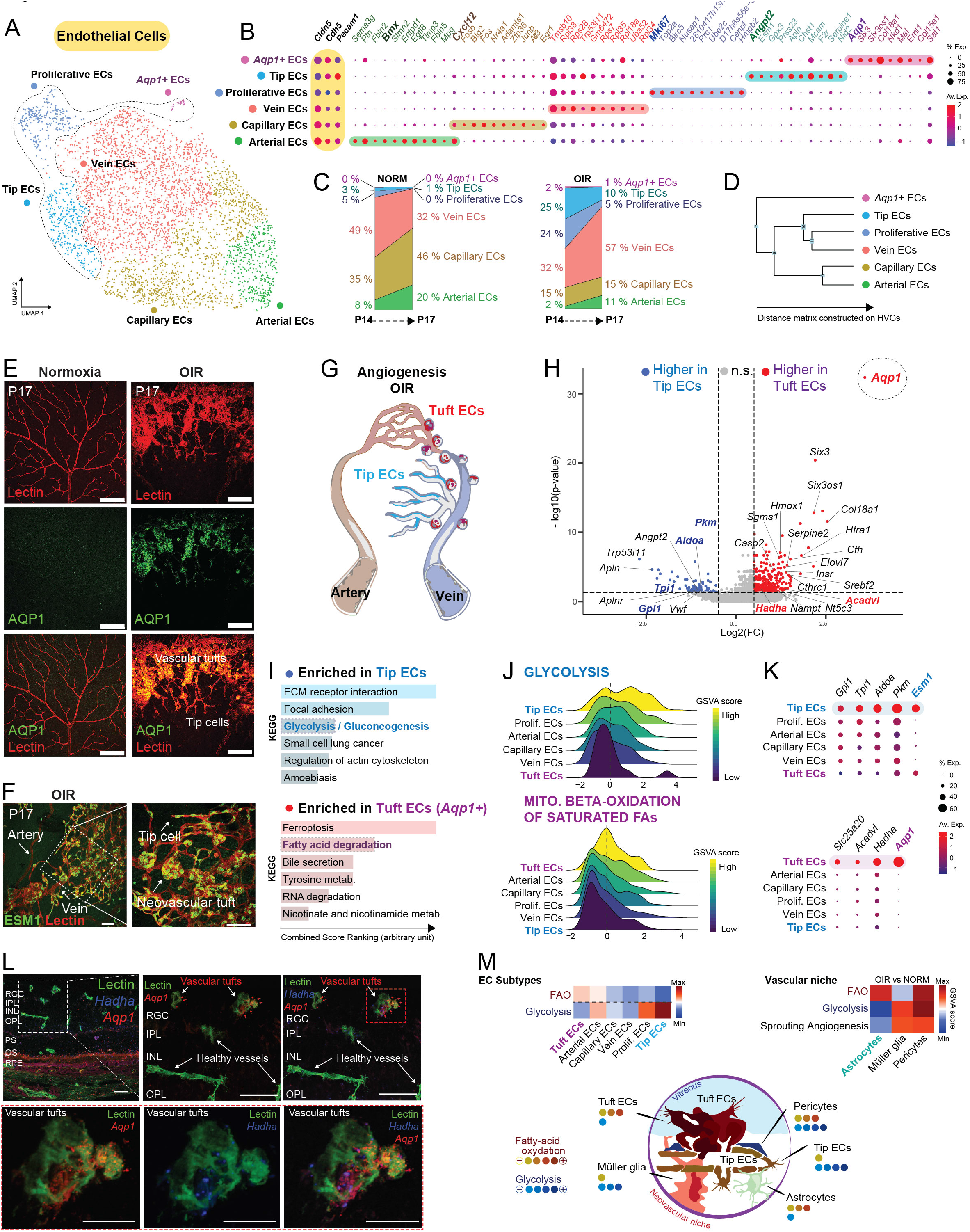
Single-cell RNAseq identifies a unique transcriptional signature for neovascular tufts. **A.** UMAP of endothelial-enriched retinal single-cell RNAseq from normoxic (n=4, 2094 cells) and OIR (n=5, 1875 cells) retinas during the neovascular phase (P14-P17) representing 6 endothelial cell subtypes identified by graph-based clustering of normalized RNA count. **B.** Dot plot of the top 10 marker genes from each endothelial subtype identified on UMAP. **C.** Representation of the percentage of cells within each endothelial subtype, calculated from the total number of endothelial cells for each condition and at each time point. **D.** Hierarchical dendrogram of endothelial subtypes using k-mean Euclidean distances constructed on high variable genes (HVGs). **E.** Immunofluorescence of retinal flat-mounts for AQP1 (green) counterstained with Lectin (red) of mice exposed to normoxia or OIR (P17). Scale: 150 μm. **F.** Immunofluorescence of P17 OIR retinal flat-mounts for ESM1 (green) and Lectin (red). Scale: 10 μm. **G.** Graphical representation of physiological (tip cells) and pathological (Tuft) angiogenesis during neovascularisation in OIR **H.** Volcano plot presenting genes differentially expressed between tuft and tip endothelial cells, derived from single-cell RNAseq data of OIR retinas. Inclusion criteria are a P-value <0.05, and an absolute log2 Fold Change (FC) > 0.5, analyzed by a non-parametric Wilcoxon rank-sum test, excluding genes expressed in fewer than 10% of cells. **I.** KEGG pathway analysis of up-regulated genes in tip or tuft EC compared to each other (combined P-value and z-score). **J.** Ridge plot of normalized GSVA score for glycolysis (Hallmark) and mitochondrial beta-oxidation of saturated fatty acids (Reactome) pathways in OIR endothelial subtypes at P17. **K.** Dot plot of the most expressed glycolytic and FAO genes from the metabolic pathways in J. **L.** *In situ* hybridization by RNAscope of P17 OIR retinal cryosection for the endothelial subtype marker *Aqp1* (red) and the FAO enzyme *Hadha* (blue) counterstained with Lectin (green). Scales: 25 μm (top panels), 10 μm (bottom panels). **M.** Heat map of GSVA scores for metabolic and angiogenic pathways in endothelial cell (EC) subtypes (left) and the vascular niche (right) comparing OIR to normoxia. Graphical model of the metabolic reliance on FAO and glycolysis of cells of the vascular unit (bottom). See also Figure S3.

Exclusively in OIR retinas, we detected a small EC cluster (1 to 2 % of total EC) defined by the specific expression of Aquaporin 1 (*Aqp1*), a water channel that controls cellular osmotic homeostasis. These unique *Aqp1*+ cells co-expressed *bona fide* EC genes (*Cdh5, Cldn5, Pecam1*) as well as vein (*Rpl18a, Tmsb10, Rps28*) and tip cell (*Esm1, Mcam, Prss23*) markers. Still, they were unique by their expression of genes involved in cell permeability (*Aqp1*) and matrix remodeling (*Col18a1, Col15a1*), amongst other pathways (Figures 3B and S3C, D). We also confirmed the presence of *Aqp1+* endothelial subclusters in publicly available scRNAseq datasets from mouse OIR retinas (P14) (Figure S3E)^28^. Hierarchical clustering based on highly variable genes showed that the *Aqp1*+ EC were the most divergent compared to other endothelial subtypes and closest to the tip cell subcluster (Figure 3D). Based on these transcriptional characteristics and the absence of *Aqp1*+ EC in normoxic retinas (devoid of tufts), we postulated that *Aqp1*+ EC might correspond anatomically to pathological neovascular tuft EC. Indeed, we localized AQP1-expressing EC to pathological tufts in OIR retinas using immunofluorescence but not in healthy normoxic vessels (Figure 3E). We also confirmed the expression of tip cell marker ESM1 in both tip and tuft EC (Figure 3F), in line with previous evidence that tuft EC might be misguided tip cells^8^. Hence, we identified a unique transcriptional signature for pathological neovascular tuft EC in the murine OIR model, characterized by AQP1 expression.

### FAO and glycolysis dichotomize neovascular tufts and physiological tip cells

We then compared the newly obtained transcriptional signature of pathological tuft EC to physiological tip EC required for regenerative angiogenesis. We identified 449 differentially expressed genes (DEGs) between tuft and tip EC (*P* value < 0.05, log2(FC) > 0.5, Figure 3G, H). Up-regulated genes in tip EC were enriched for extracellular matrix-receptor interaction, focal adhesion, and glycolysis pathways. In contrast, tuft EC genes were significantly enriched for ferroptosis, fatty-acid degradation, tyrosine, and nicotinamide metabolism pathways (Figure 3I). To better visualize the metabolic dichotomy between tuft and tip EC, we performed a gene set variation analysis (GSVA) to rank EC subtypes based on their relative expression of genes implicated in glycolysis and FAO pathways (Figure 3J). In OIR, tip EC were highly glycolytic and relied less on FAO (P17). Inversely, tuft EC relied preferentially on FAO and least on glycolysis (P17).

Focusing on the most abundant FAO and glycolysis transcripts, tip cells showed greater expression of essential glycolic genes (*Pkm, Aldoa, Tpi1, Gpi1*), particularly at P17, while tuft EC expressed higher levels of FAO genes (*Hadha, Acadvl*) (Figures 3K). Notably, tuft EC strongly expressed *Hadha* (Hydroxyacyl-CoA Dehydrogenase Trifunctional Multienzyme Complex Subunit Alpha), which encodes a subunit of the mitochondrial trifunctional protein essential for FAO. Importantly, *Hadha* expression colocalized with pathological *Aqp1*+ tufts by RNAscope *in situ* hybridization on retinal cryosections (Figure 3L). Thus, we discerned distinct transcriptional metabolic identities for healthy and diseased EC in murine PR, favoring glycolysis in the case of tip EC and FAO in the case of pathological tuft EC (Figure 3M, left). This metabolic dichotomy was also evident in the vascular niche, with astrocytes highly expressing FAO genes, while Müller cells were more glycolytic in OIR (Figures 3M, right). Importantly, we offer an endothelial atlas from the ischemic phase of OIR (P14/P17), enabling deeper insights into the divergent dynamics separating pathological and physiological angiogenesis.

### Targeting FAO curbs pathological neovascular tufts

We then explored the role of FAO in neovascular tuft formation. We utilized etomoxir to inhibit FAO, blocking CPT1a, essential for transporting lipids into mitochondria^16^ (Figure 4A). Etomoxir reduced pathological neovascularization (NV) and vaso-obliteration (VO) in WT retinas exposed to OIR (Figure 4B), in line with a contribution of FAO to neovascular tuft formation. However, etomoxir caused substantial toxicity and fatalities during the hypoxic phase of OIR (P12 to P17), highlighting the vital role of FAO metabolism systemically. We, therefore, sought to identify a more targeted metabolic regulator capable of yielding therapeutic gains without harmful systemic side effects.

**Figure 4.**
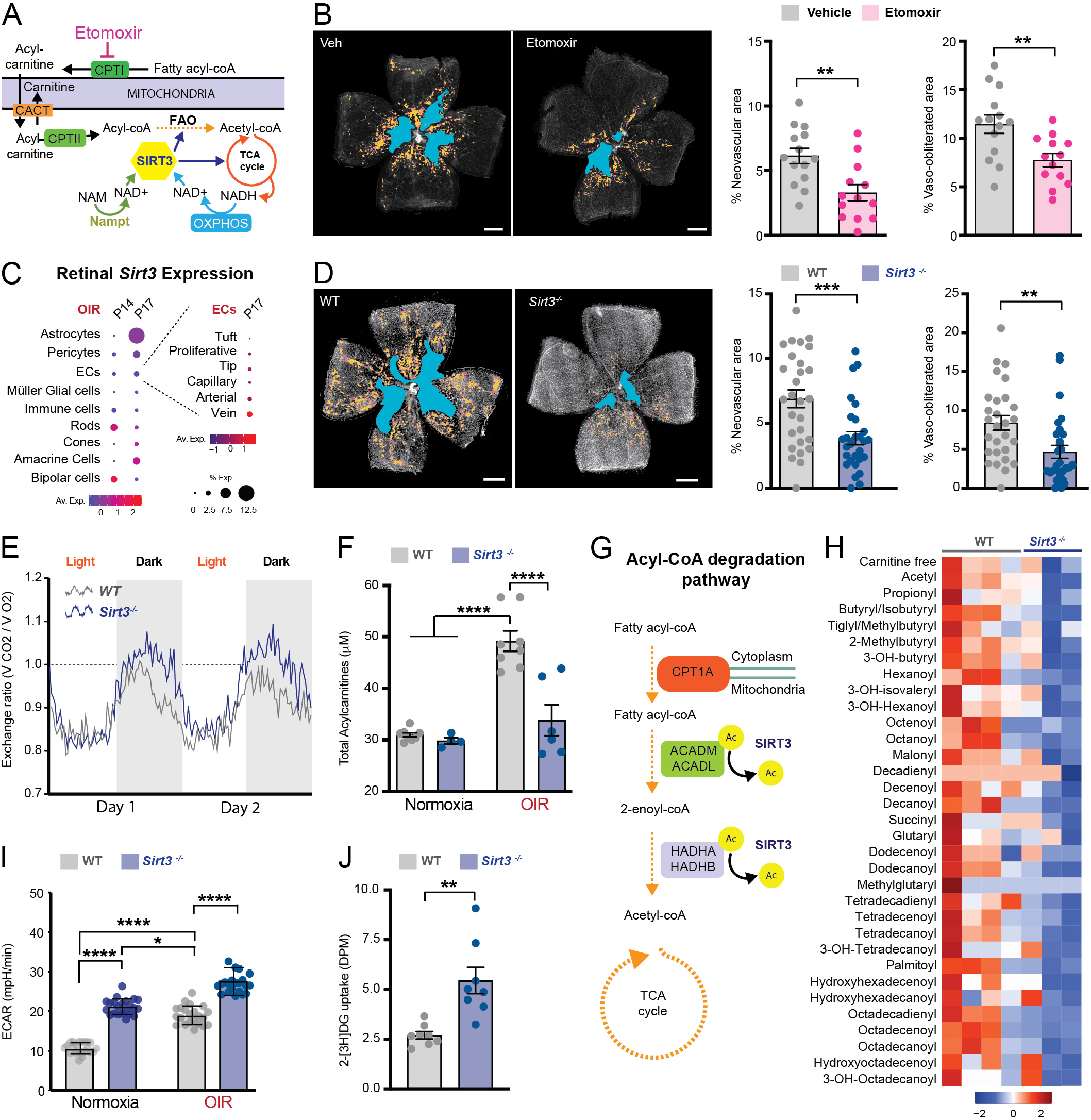
Targeting FAO curbs pathological neovascular tufts. **A**. Graphical representation of Sirtuin-3 regulation of mitochondrial fatty acid oxidation (FAO). Etomoxir inhibits CPT1A which blocks FAO. **B.** Lectin-stained retinal flat-mount of P17WT mice exposed to the OIR model and treated with vehicle (n= 14 retinas) or etomoxir (n= 13 retinas) from P12 to P17; vaso-obliterated (VO, blue) and neovascular (NV, yellow) areas are highlighted. Bar graph of VO and NV areas relative to the total retinal area. Scale: 500 μm. **C.** Dot plot of retinal Sirt3 expression in OIR (P14 and P17) and endothelial cell (EC) subtypes (P17). Sirt3 expression only increased significantly in astrocytes (12.5% of cells, P= 0.008). **D.** Lectin-stained retinal flat-mount of WT and *Sirt3^-/-^* mice (n=28retinas each) exposed to the OIR model (P17); vaso-obliterated (VO, blue) and neovascular (NV, yellow) areas are highlighted. Bar graph shows the VO and NV areas relative to the total retinal area. Scale: 500 μm. **E.** Metabolic cages were used to measure the O2 consumption (VO2) and CO2 production (VCO2) of *Sirt3*^-/-^ and WT mice (n= 3 mice/group). Respiratory Exchange ratio (RER) of ≥1 is indicative that carbohydrates are the predominant fuel source. **F.** Bar graph of total acylcarnitine levels measured by LC/MS/MS of normoxic and OIR retinas from WT (NORM n=8 retinas, OIR n = 8 retinas) and *Sirt3^-/-^* mice (NORM n = 4 retinas, OIR n = 6 retinas) at P17. **G.** Graphical representation of fatty acyl-coA degradation pathway. **H.** Heatmap of fatty acylcarnitine metabolites in OIR WT (n = 4 mice) and *Sirt3^-/-^* (n=3 mice) from acylcarnitine-targeted metabolomics analysis of pooled retinas (2 retinas per pool). **I.** Basal extracellular acidification rate (ECAR) measured by Seahorse analyzer on retinal explants exposed to normoxia (Norm) or OIR from WT (n=40 retinal punches, from 10 retinas) and *Sirt3^-/-^* retinas (n=28 retinal punches, from 7 retinas) averaged over 5 basal time points each. **J.** *In vivo* glucose uptake in WT (n = 8 retinas) and *Sirt3^-/-^* (n = 8 retinas) retinas following OIR. Retinal radioactivity counts of 2-[^3^H] deoxyglucose (DG) tracer relative to the total injected dose show glucose uptake by retinal cells at P14. DPM: Disintegration per minute. Statistical significance was established using two-tailed Student’s *t*-test (**B, D, J**) and ANOVA with Tukey post hoc analysis (**F, I**). **P<0.01, ***P<0.001, ****P<0.0001 See also Figure S4.

Mitochondrial SIRT3 was an interesting candidate due to its central role in regulating cellular energy homeostasis without entirely inhibiting FAO. SIRT3 enhances the activity of essential enzymes of FAO^20^, the Krebs cycle^29,30^, and the respiratory chain^31^ (Figure 4A). SIRT3 directly deacetylates HADHA and ACADVL^32^ (Figure S4A), two FAO enzymes predominantly enriched in the neovascular unit (Figure 2E). During OIR, *Sirt3* expression increased significantly in astrocytes (12.5% of cells, P = 0.008), whereas its expression was limited in EC (2%) and absent in tuft EC by scRNAseq (Figure 4C). Accordingly, conditional EC deletion of *Sirt3* (*Cdh5-CreERT2, Sirt3^lox/lox^*) did not impact the vascular phenotype of transgenic mice exposed to OIR (Figure S4B-D). Interestingly, global *Sirt3* deletion (*Sirt3^-/-^*) markedly reduced pathological neovascular tuft (NV) and vaso-obliteration (VO) compared to wild-type (WT) retinas (Figure 4D), akin to the effects of etomoxir but without increased mortality or apparent deleterious consequences. Hence, *Sirt3* depletion of the neurovascular niche, but not EC alone, reduced pathological angiogenesis.

Next, we examined the metabolic effects of *Sirt3* depletion. Using metabolic cages, *Sirt3^-/-^* mice displayed a higher exchange ratio (VCO2/VO2) during their active nocturnal phase, indicating a preference for carbohydrate metabolism over lipid reliance (Figure 4E). Their acyl-carnitine profiles revealed that *Sirt3* deletion prevented the increase in total acylcarnitines noted in WT retinas during OIR (Figure 4F). Indeed, reduced levels of fatty acylcarnitine metabolites were measured in mutant retinas compared to WT during OIR (Figure 4G, H), underscoring the importance of SIRT3 in mitochondrial FAO. To assess their glycolytic rate, we measured the extracellular acidification rate (ECAR) of retinal explants using a Seahorse analyzer. *Sirt3^-/-^* retinas exhibited higher basal ECARs, further rising under OIR conditions, suggesting an enhanced glycolytic rate (Figure 4I). *In vivo,* we also observed increased retinal glucose uptakes in *Sirt3^-/-^* retinas compared to WT, using trace amounts of radioactively labeled 2-deoxyglucose (DG) injected in mice exposed to OIR (Figure 4J); 2-DG is a glucose analog that is not metabolized by the retina. Hence, a metabolic shift from FAO to glycolysis in *Sirt3^-/-^* retinas was associated with fewer neovascular tuft formation.

### *Sirt3* depletion alters the metabolic and angiogenic landscape of the neovascular niche

To define the transcriptional effects of *Sirt3* deletion during pathological angiogenesis, we repeated a single-cell RNAseq of EC-enriched retinal cell suspensions. The enrichment strategy improved EC yields, but other retinal cell types were also detected (Figure 5A, top). To identify which retinal cell types were most impacted by the global deletion of *Sirt3* at a transcriptional level, we employed a machine-learning approach using a Random Forest Classifier termed Augur^33^. Based on Augur results, EC, and macroglial cells, comprising Müller cells and astrocytes, were the most impacted cell types in our model (Figure 5A, bottom). In *Sirt3* depleted retinas, the EC composition transitioned rapidly from a more proliferative neovascular profile at P14 compared to WT (tuft, tip, and proliferative EC: 43.7% vs. 38.2%) to a more quiescent EC composition at P17 (16.1% vs 23.7%), resembling the normoxic retina (Figure 5B, C); tuft EC were barely detected (0.1 % vs 0.7%).

**Figure 5.**
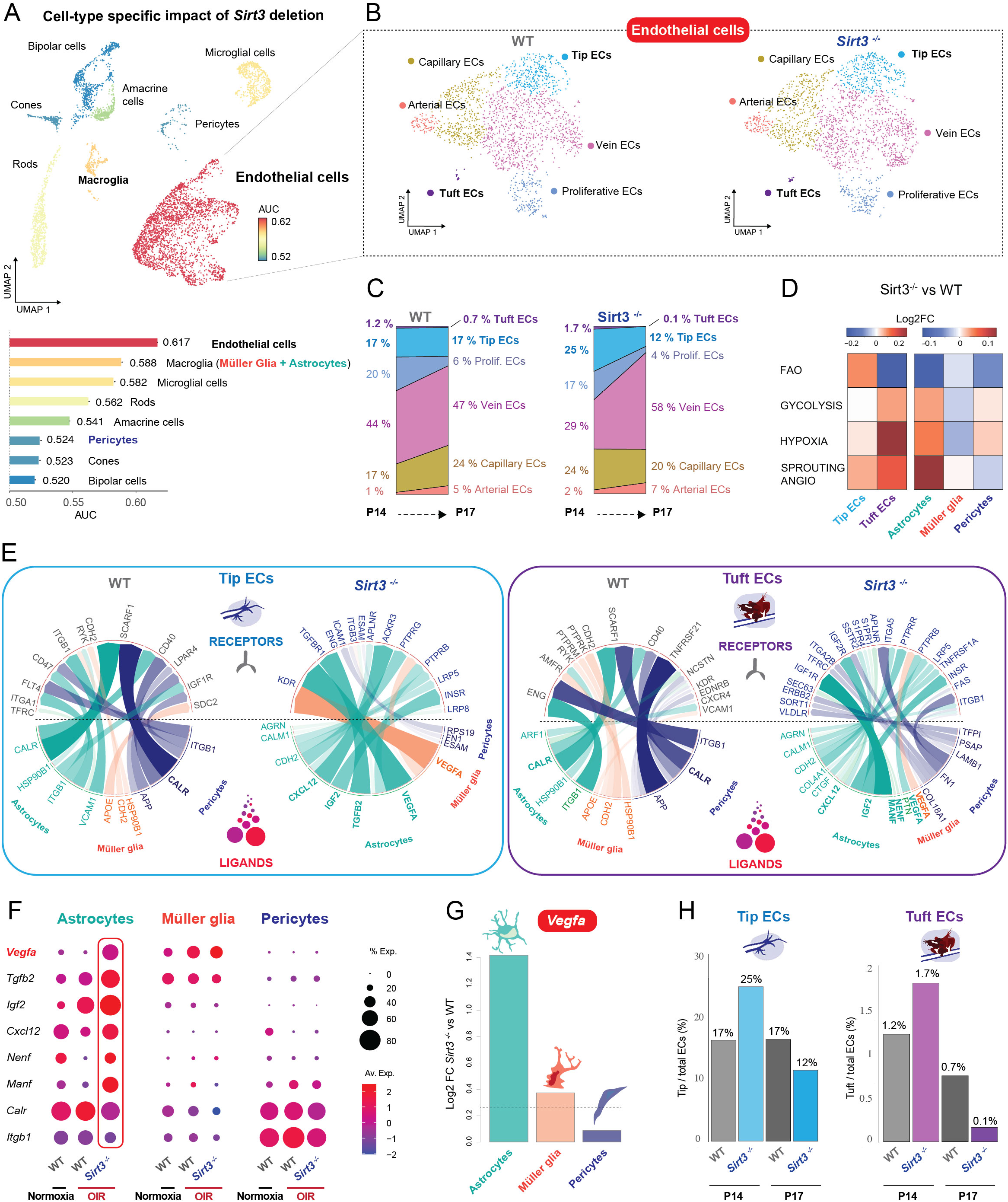
*Sirt3* depletion alters the metabolic and angiogenic landscape of the neovascular niche. **A.** UMAP and lollipop plot of cell type prioritization score from AUGUR analysis on single-cell RNAseq analysis of WT (n=5, 4047 cells) and *Sirt3*^-/-^ (n=5, 3639 cells) retinas during the neovascularisation phase of OIR (P14-P17). AUC = area under the curve. **B.** UMAP of endothelial subclusters from WT (n=5, 1875 cells) and *Sirt3*^-/-^ (n=5, 1949 cells) retina during the neovascularisation period (P14-P17) representing the 6 EC subtypes identified by graph-based clustering of normalized RNA count. **C.** Bar graphs represent the percent of cells in each endothelial subtype out of the total number of EC per genotype and time point. **D.** Heatmap representing the log2 fold change (FC) of normalized GSVA score for metabolic and angiogenic pathways between WT and *Sirt3^-/-^* tip and tuft EC, and neuroretinal cells from scRNAseq data of OIR retinas (P14 and P17). **E.** Circos Plot from NicheNet analysis showing the differential communication between WT and *Sirt3^-/-^* neurovascular ligands and tip and tuft EC receptors in OIR retinas based on scRNAseq data (P14/P17 combined). Selected ligands (most potent) were differentially expressed between WT and *Sirt3*^-/-^ neurovascular cells (log2FC>0.25, % expression > 40%). Selected receptors were differentially expressed between WT and *Sirt3^-/-^* tip (left) and tuft (right) EC (log2FC>0.25, % expression > 10%). **F.** Dot plot representing the differential expression of selected ligands from WT and *Sirt3*^-/-^ retinal neurovascular cells based on NicheNet analysis. **G.** Bar graph depicting Log2 FC of *Vegfa* expression in *Sirt3^-/-^* vs. WT vascular unit cells. **H.** Percent of tuft EC (WT = 16 cells, *Sirt3*^-/-^ = 15 cells) and tip cells (WT = 321 cells, *Sirt3*^-/-^ = 327 cells) out of the total number of EC per genotype and time point. See also Figure S5.

Reduced pathological angiogenesis in *Sirt3*-depleted retinas (Figure 4D) was associated with a metabolic shift from FAO to glycolysis within the vascular niche, particularly in tuft EC and astrocytes (Figure 5D). In turn, this change in metabolism altered the interactome of the vascular niche with tip and tuft EC (Figure 5E), dampening inflammatory and proliferative signals observed in WT retinas during OIR (Figure S5A-D). Notably, astrocytes of *Sirt3*-depleted retinas expressed more VEGF (5-fold increase) and other important angiogenic factors (Figure 5F, G) to levels comparable during retinal development (P6-P10) (Figure S5E-G). Astrocytes classically form a physical path and secrete VEGF, enabling physiological vascular growth^3,6^. In OIR, the shift in the vascular niche metabolism and the higher VEGF secretion from astrocytes were associated with more tip and tuft EC in *Sirt3^-/-^* retinas at P14 (Figure 5H). At this early stage in retinal revascularization (P14), neovascular cells are not yet fully committed, and tufts are barely visible histologically. Surprisingly, fewer tuft EC were observed at P17 in *Sirt3^-/-^* retinas, raising the possibility that early tuft EC fate was altered as a result of their microenvironment.

### *Sirt3* deletion reprograms early neovascular tufts for regenerative angiogenesis

Given that EC behavior adapts to their environment, we explored whether the angiogenic shift noted in the *Sirt3^-/-^* neovascular niche could prompt tuft EC to adopt a tip-like endothelial fate. Indeed, despite an early increase in tuft EC at P14, the ratio of tip cells to tuft EC significantly increased in *Sirt3^-/-^*retinas at P17 (Figure 6A), suggesting a transition from pathological to physiological angiogenesis in the mutant retinas. Moreover, analysis of transcriptional trajectory (using Monocle 3) showed that in contrast to WT, tuft EC were still connected to tip cells in *Sirt3^-/-^*retinas (Figure 6B), raising the possibility that they could be reprogrammed to take on a physiological fate.

**Figure 6.**
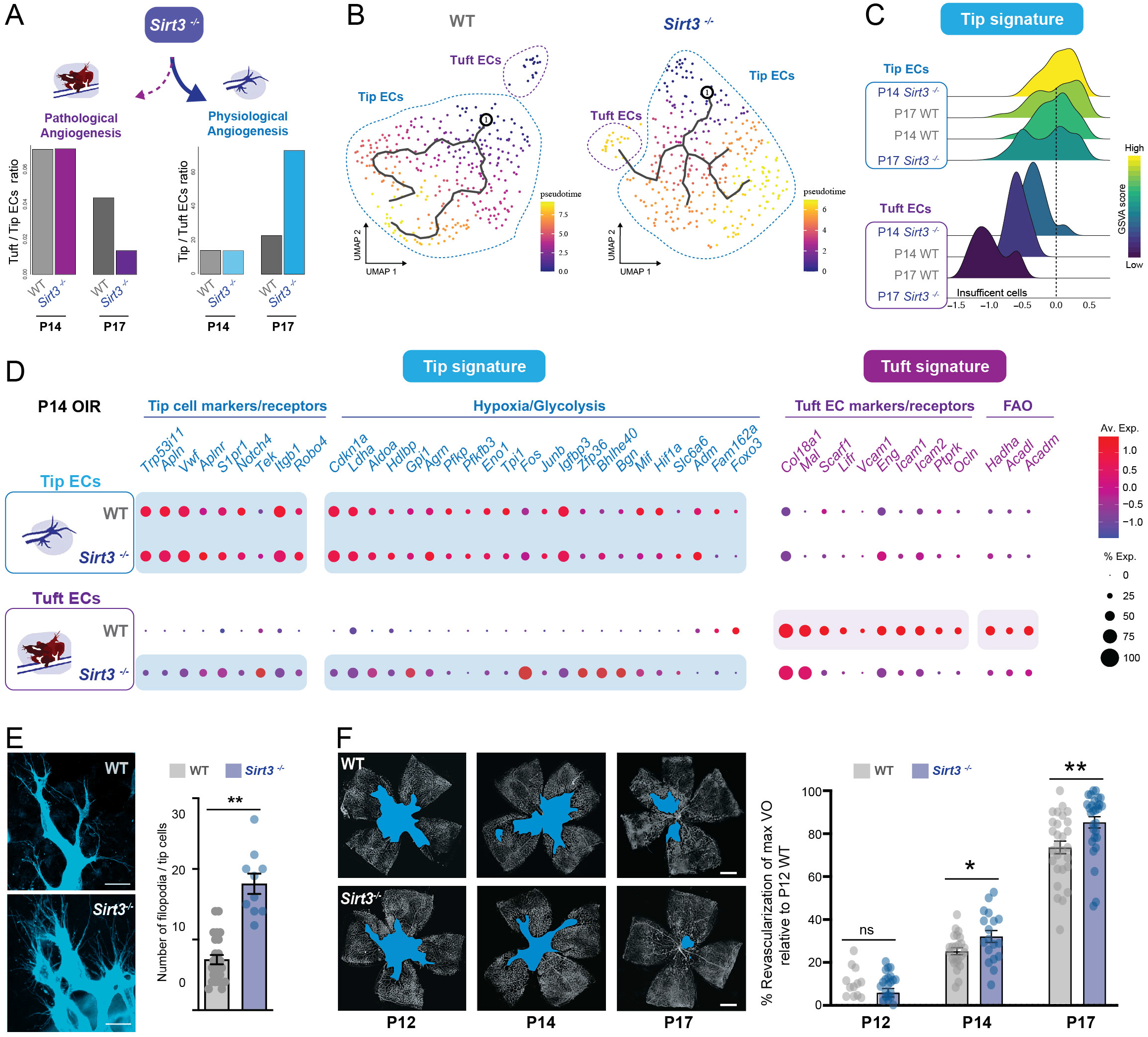
*Sirt3* deletion reprograms early neovascular tufts for regenerative angiogenesis. **A.** Bar plot representing the ratio of tip to tuft EC (left) and inversely (right) during the neovascularisation phase (P14-P17) in WT and *Sirt3*^-/-^ OIR retinas. **B.** UMAP of single-cell RNAseq from WT and *Sirt3*^-/-^ neovascular cells (tip cells and tuft EC) during the neovascularisation period (P14-P17), showing the connectivity between cells through a trajectory analysis performed with Monocle 3. **C.** Ridge plot of tip and tuft EC from OIR WT and *Sirt3*^-/-^ retinas at P14 and P17 representing the normalized GSVA score for the Physiological angiogenesis gene set, as previously defined from the Tip cell signature (Figure 3H). **D.** Dot plot of gene expression in tip and tuft EC at P14 from OIR WT and *Sirt3*^-/-^ retinas focusing on pathways previously identified as related to tip and tuft EC. **E.** Representative images of tip cells forming motile filopodia in WT and *Sirt3*^-/-^ OIR retinas at P14; stained with lectin. Quantification of the number of filopodia per tip cells in WT (n=23 tip cells) and *Sirt3*^-/-^ (n=10 tip cells) retinas exposed to OIR. **F.** Percent retinal revascularization of max VO relative to P12 WT. We compared *Sirt3*^-/-^ (P12 n = 26 retinas, P14 n = 18 retinas, P17 n = 30 retinas) to WT retinas at each timepoints (P12 n = 26 retinas, P14 n = 24 retinas, P17 n = 28 retinas). Statistical significance was determined using two-tailed Student’s t-tests (E, F). *P<0.05, **P<0.01, ****P<0.0001. See also Figure S6.

Considering that FAO distinctly characterizes tuft EC (Figure 3), we examined the expression of FAO genes in tuft EC of *Sirt3^-/-^* retinas. Using GSVA and differential quantitative set analysis for gene expression (QuSAGE^34^), we identified a marked reduction in FAO pathways in *Sirt3^-/-^* tuft EC (Figure S6A), primarily due to reduced gene expression of *Acadl, Acadm,* and *Hadha* (Figure S6B). Given the predisposition of tuft EC to proliferate, possibly aided by FAO^16^, we compared the expression of cell cycle genes coding for proliferation (G2/M) or cell cycle arrest (G1) in both tuft and tip EC. We observed that, at P14, tuft EC in *Sirt3^-/-^*retinas were less proliferative than WT (30% vs. 80%), akin to the reduced proliferation index of migratory tip cells (∼30%, Figure S6C). Hence, *Sirt3* deletion prevented tuft EC from acquiring defining traits of tuft identity.

Finally, we investigated if the initial reduction in FAO and proliferation of tuft EC in *Sirt3^-/-^* retinas might correlate with a comprehensive transcriptional shift towards a tip-cell-like identity. Using GSVA and a pre-established tip cell gene signature (Figure 3H), we found that early tuft EC (P14) exhibited a transcriptional profile resembling physiological tip cells, more so than their WT counterparts (Figure 6C). We then analyzed pathways that define tip cell identity at P14 (Figure 3I), a critical time point for EC fate specification in OIR. Tip cells in WT and *Sirt3^-/-^*retinas shared analogous expression profiles (Figure 6D). However, tuft EC of *Sirt3^-/-^*retina began expressing tip cell markers, focal adhesion, hypoxia-regulated, and glycolytic genes (Figure 6D) to levels comparable to tip cells. The findings were corroborated by the observed elevation of hypoxia, glycolysis, and sprouting angiogenesis pathways in *Sirt3^-/-^* retinas, in contrast to WT (Figure 5D and S6D). These pathways are characteristic transcriptional traits of tip cells^35^. Correspondingly, *Sirt3* deletion markedly increased the number of early tip cell filopodia (P14; Figure 6E) and accelerated revascularization of the ischemic neuroretina in OIR at P14 and P17 without significantly affecting maximal vaso-obliteration at P12 (Figure 6F). Hence, *Sirt3* deletion within the neovascular niche shifted the transcriptional identity of tuft EC to resemble tip cells, thereby accelerating the physiological vascular regeneration of the ischemic neuroretina in PR.

## DISCUSSION

Recognizing the unique attributes of healthy and diseased blood vessels and their microenvironment could pave the way for more targeted and efficacious therapies against vascular diseases. This study uncovered a distinct transcriptional signature for pathological tuft EC, a defining feature of proliferative retinopathy (PR) and a leading cause of blindness. Unlike healthy tip cells, these pathological EC exhibited a preferential metabolic dependency on FAO. Notably, shifting the metabolic program of the vascular niche from FAO to glycolysis in *Sirt3^-/-^* retinas reduced pathological neovascular tufts and promoted healthy retinal revascularization.

Pathological neovascular tufts are portrayed as misdirected tip cells invading the vitreous^8^ that are more proliferative and leaky, resulting in retinal edema^36^. Through single-cell transcriptomics, we found a novel subset of EC that, while displaying multiple tip cell markers such as ESM1, were less prone to migrate and more inclined to proliferate. The new EC sub-cluster expressed high levels of aquaporin 1 (AQP1) that we localized to pathological neovascular tufts. AQP1 is a water channel pivotal for blood-retinal barrier permeability^37,38^ and could contribute to vascular leakage associated with PR^37,38^. In tumor models, *Aqp1* directly contributes to EC migration, sprouting, and tubulogenesis^39^, implying a direct role in pathological angiogenesis^40^. Hence, AQP1 uniquely labels pathological tuft EC in PR and could be a new therapeutic target to curb retinal edema and pathological neovascularization in proliferative vascular diseases.

Using their distinct transcriptional signature, we showed that unique metabolic profiles defined tip and tuft EC. In both human neonates and mice, aberrant vascular growth is ushered by an increase in the neuroretina’s metabolic demands, pointing to the pivotal role that retinal metabolism plays in determining EC fate^4^. Unbiasedly comparing the most enriched transcriptional pathways of tip and tuft EC revealed divergent metabolic preferences: tip cells favored glycolysis, whereas tuft cells leaned towards FAO. Notably, glycolysis is less dependent on oxygen for energy production and cellular growth, a phenomenon known as the Warburg effect, while FAO necessitates mitochondrial respiration^41^. This metabolic specialization aligns anatomically with the location of tip EC at the leading edge of the oxygen-poor, avascular neuroretina. In contrast, neovascular tufts are found at the confluence of venous capillaries^42^, an area richer in oxygen. FAO is a fuel source for the neuroretina^43^ and serves as a biosynthetic pathway promoting EC proliferation^16^. Unlike glycolysis, FAO is not associated with increased migration typical of tip cells^12^. Hence, distinct metabolic pathways define the behavior of glycolytic tip cells, essential to the regenerative process, in contrast to misguided yet actively proliferating tuft EC that rely on FAO. Crucially, these distinct metabolic profiles for healthy and pathological angiogenesis become apparent at specific stages of disease development, presenting a potential therapeutic window for targeted interventions.

Localization of the metabolic and angiogenic signals shapes regenerative angiogenesis. *Sirt3* expression was only significantly increased in astrocytes in mouse PR and was not detected in tuft EC. In the absence of *Sirt3*, astrocytes exhibited a pronounced metabolic shift towards glycolysis and elevated angiogenic signaling, with a 5-fold increase in Vegfa expression, comparable to levels seen in the developing retina (P6). Given that astrocytes act as a scaffold to guide migrating EC during retinal development, they could provide localized signals to guide beneficial revascularization in PR. PR treatments aim to lower VEGF concentrations after neovascular tuft formation has taken place, irrespective of its source. Although these approaches yield initial improvements in macular edema and visual acuity^44^, primarily within the first six months, the benefits wane over time^45^ and are associated with anticipated neuronal toxicities^46^. Our findings highlight the importance of the cellular source of growth factors, including VEGF, to guide healthy retinal revascularization, not merely its concentration. VEGF has multiple neuroprotective roles that could be leveraged to enhance neuronal function^47^. Therefore, modifying the metabolic topography of the vascular niche, especially in astrocytes, could help steer EC toward a more regenerative path.

Maintaining lower oxygen saturation levels is the mainstay of ROP prevention^48^ and likely contributes to EC reprogramming. Within limits, however, since more severe hypoxia, although beneficial to ROP, is associated with increased mortality^49^. Hypoxia blocks mitochondrial respiration, forcing tissues to rely on glycolysis instead^50,51^, as we observed in SIRT3-depleted mice. In mouse models of mitochondrial diseases, exposure to chronic hypoxia is protective by diverting energy metabolism away from mitochondrial oxidative phosphorylation^50^. In the OIR model, pharmacological stabilization of hypoxia-inducible factor (HIF) also shifts the retina towards aerobic glycolysis, improving retinopathy^52^. Similarly, Sirt3 null mice shifted their metabolism from FAO towards glycolysis^53^, recreating a pseudo-hypoxic state that could pre-condition mice for ischemic stress in PR. Thus, if administered locally in the eye, targeting SIRT3 could favor a regenerative phenotype while avoiding some of the harmful consequences of severe hypoxia. Then again, SIRT3 was reported to have protective antioxidant roles in photoreceptors^54^. Fortunately, SIRT3 was not significantly expressed in photoreceptors during PR, and *Sirt3* depletion alone did not impact vision in other models, suggesting protective effects of other mitochondrial sirtuins, such as SIRT5^55,56^.

Our current therapies target pathological neovessels that emerge in the later phases of PR. In contrast, our research aims to enable early physiological vascular regeneration to minimize neuroretinal ischemia. Targeting SIRT3 shifted the metabolic environment of the vascular niche, steering EC fate towards a regenerative vascular phenotype. This work expands our understanding of the distinct metabolic and transcriptomic landscapes of healthy and diseased vessels and their microenvironment, offering insights for developing targeted therapies against vascular diseases^57,58^. Additionally, our findings have implications for other neuro-ischemic conditions like stroke, where enhancing regenerative angiogenesis could preserve neuronal function^59^.

## ACKNOWLEDGMENTS

We thank Marie-Josée Lacombe for her technical assistance. We thank Elke Küster-Schöck and the Platform for Imaging by Microscopy of CHU Sainte-Justine Research Center (CHUSJRC), supported by Leica Microsystems and funded by CHUSJRC, the Quebec government, CHUSJ Foundation and Canada Foundation for Innovation and the Vision Health Research Network for funding of the Single-cell Genomics Analysis Platform. GC was supported by a MITACS Elevate fellowship. J.-S.J. was supported by the Burroughs Wellcome Fund, the Canadian Institute of Health Research (CIHR; 390615, 479607), the National Sciences and Engineering Research Council of Canada (NSERC; 06743), and the Fonds de Recherche du Québec–Santé (FRQS). PS was supported by CIHR (353770), Heart & Stroke Foundation Canada (G-16-00014658), Foundation Fighting Blindness Canada, and the Canadian Diabetes Association (DI-3-18-5444-PS).

## AUTHOR CONTRIBUTIONS

G.C., S.P. and J-S.J. conceived and designed all experiments and wrote the manuscript; F.R. and P.S. collected human samples; C.B.C. generated the metabolomic data of human samples; G.C analyzed the human metabolomics data; B.M. and S.P. performed the carnitine quantification of mouse retinal sample and G.C. analyzed the data; S.P. and P.G. performed the Seahorse experiments; G.C., F.W., S.L. and G.A. generated single-cell RNAseq (Drop-seq) data; G.C., A.S. and M.X.C. analyzed single-cell RNAseq data; S.P., J.S.K., N.R.H., T.A. and C.B. performed oxygen-induced retinopathy experiments and phenotypic quantification; S.P, J.S.K., E.H. and N.R.H. performed *in vivo* and *in vitro* metabolic experiments; J.C.R. performed immunohistochemistry experiments; E.H. performed RNAscope experiments; S.C.G. provided graphic design expertise, A.D., L.E.H.S. and P.S. provided valuable insight regarding retinal vascular biology; all authors analyzed the data.

## INCLUSION AND DIVERSITY

We support inclusive, diverse, and equitable conduct of research.

## Nonstandard Abbreviations and Acronyms

DR: Diabetic retinopathy
PR: Proliferative retinopathies
PDR: Proliferative diabetic retinopathies
NV: Neovascularization
OIR: Oxygen-induced retinopathy
VO: Vaso-obliteration
EC: Endothelial cells
NVT: Neovascular tuft
P14: Post-natal day 14
P17: Post-natal day 17
FAO: Fatty acid beta-oxidation
2-DG: 2-deoxyglucose
ERG: Electroretinography
GSVA: Gene set variation analysis
DEGs: Differentially expressed genes
*Sirt3*: Sirtuin-3
*Cpt1*: Carnitine palmitoyltransferase 1
*Acadl*: Acyl-CoA dehydrogenase
*Acadvl*: Very long-chain specific acyl-CoA dehydrogenase
*Hadh*: Hydroxyacyl-Coenzyme A dehydrogenase
*Hadha*: Hydroxyacyl-CoA Dehydrogenase Trifunctional Multienzyme Complex Subunit Alpha
*Hadhb*: Hydroxyacyl-CoA Dehydrogenase Trifunctional Multienzyme Complex Subunit Beta
*Pecam1*: Platelet and Endothelial Cell Adhesion Molecule 1
*Esm1*: Endothelial Cell-Specific Molecule 1
*Aqp1*: Aquaporin 1
*Col18a1*: Collagen Type XVIII Alpha 1 Chain
*Col15a1*: Collagen Type XV Alpha 1 Chain
*Pkm*: Pyruvate Kinase M1/2
*Aldoa*: Aldolase, Fructose-Bisphosphate A
*Tpi1*: Triosephosphate Isomerase 1
*Gpi1*: Glucose-6-Phosphate Isomerase

## STAR METHODS

### RESOURCE AVAILABILITY

#### Lead contact

Further information and requests for reagents should be directed to the LEAD CONTACT, Jean-Sebastien Joyal.

#### Material availability

All unique/stable reagents generated in this study are available from the lead contact with a completed Material Transfer Agreement.

#### Data and code availability

Custom-written scripts used in this study are available from the corresponding author upon reasonable request. Any additional information required to reanalyze data reported in this paper is available from the lead contact.

### EXPERIMENTAL MODEL AND SUBJECT DETAILS

#### Human Subjects

The study conforms to the tenets of the Declaration of Helsinki, and approval of the human clinical protocol and informed consent were obtained from the Maisonneuve-Rosemont Hospital Ethics Committee (CER: 10059). All patients previously diagnosed with proliferative diabetic retinopathy or epiretinal membrane were followed clinically, and surgery was performed when indicated by ’standard-of-care’ guidelines by a single vitreoretinal surgeon (Table S1). Undiluted vitreous samples were aspirated in the region adjacent to neovascular tufts or epiretinal membrane (control) and were frozen on dry ice immediately after biopsy and stored at –80°C.

#### Animal care

Animal procedures were performed in compliance with the Animal Care Committee of CHU Sainte-Justine, following the principles of the Guide for the Care and Use of Experimental Animals developed by the Canadian Council on Animal Care. All *in vivo* work adhered to the Association for Research in Vision and Ophthalmology Statement for the Use of Animals in Ophthalmic and Vision Research. S129 (#002448), mutant mice with *Sirt3* deletion (129-*Sirt3^tm^*^1^*^.1Fwa^*/J, #012755), CAG-tdTomato reporter mice (#007914), and *Sirt3^lox/lox^* mice (#031201) were obtained from The Jackson Laboratory. Cdh5-CreERT2 mice were kindly provided by Dr. Ralf Adams (Max Planck Institute, Germany). All colonies were maintained and bred in identical lighting conditions at the CHU Sainte-Justine mice facility. Pups weighing less than 6 grams or more than 8 grams on post-natal (P) day 14 or 17 (P14, P17) were excluded. Both littermate females and males were used. All animals were sacrificed by lethal intraperitoneal (i.p.) injections of pentobarbital at P12, P14, or P17 for analyses.

#### Oxygen-induced retinopathy model

Mice pups and their adoptive lactating S129 mother were exposed to 75% oxygen from P7 to P12, as first described^1^. Mice were returned to room air at P12 until used for experimentation. Littermate controls, as well as age and weight-matched mice, were injected intraperitoneally twice daily with etomoxir (27 mg/kg per day) or vehicle (0.9% saline) from P12 to P17 and euthanized at P17. 4-OH tamoxifen (5 μg) was injected intraperitoneally daily for 3 days (P12-P14) and euthanized at P17.

### METHOD DETAILS

#### Human vitreous metabolomics

Human vitreous metabolite extracts were analyzed using two liquid chromatography-tandem mass spectrometry (LC-MS) methods to measure polar metabolites, as described previously^2^. From vitreous samples, negative ion mode profiling samples were prepared from 30 µl extracted with 120 µl of 80% methanol containing inosine-15N4, thymine-d4, and glycocholate-d4 internal standards (Cambridge Isotope Laboratories, MA) while samples for positive ion mode profiling were prepared by extracting 10 µL with 90 µl of acetonitrile/methanol/formic acid (74.9:24.9:0.2 v:v:v) containing valine-d8 and phenylalanine-d8 internal standards (Cambridge Isotope Laboratories, MA). Negative ion mode data were acquired by injecting extracts (10 µL) onto a 150 x 2.0 mm Luna NH2 column (Phenomenex, CA). The column was eluted at a flow rate of 400 µl/min with initial conditions of 10% mobile phase A (20 mM ammonium acetate and 20 mM ammonium hydroxide in water) and 90% mobile phase B (10 mM ammonium hydroxide in 75:25 v/v acetonitrile/methanol) followed by a 10 min linear gradient to 100% mobile phase A. MS analyses were carried out using electrospray ionization in the negative ion mode using full scan analysis over m/z 70-750 at 70,000 resolution and 3 Hz data acquisition rate. Additional MS settings were ion spray voltage, -3.0 kV; capillary temperature, 350°C; probe heater temperature, 325 °C; sheath gas, 55; auxiliary gas, 10; and S-lens RF level 50. Positive ion mode data were acquired by injecting extracts (10 µl) onto a 150 x 2 mm, 3 µm Atlantis HILIC column (Waters, MA). The column was eluted isocratically at a flow rate of 250 µl/min with 5% mobile phase A (10 mM ammonium formate and 0.1% formic acid in water) for 0.5 minutes, followed by a linear gradient to 40% mobile phase B (acetonitrile with 0.1% formic acid) over 10 minutes. MS analyses were carried out using electrospray ionization in the positive ion mode using full scan analysis over 70-800 m/z at 70,000 resolution and 3 Hz data acquisition rate. Other MS settings were: sheath gas 40, sweep gas 2, spray voltage 3.5 kV, capillary temperature 350°C, S-lens RF 40, heater temperature 300°C, microscans 1, automatic gain control target 1e6, and maximum ion time 250 ms. Raw data were processed using TraceFinder 3.3 and 4.1 software (Thermo Scientific, MA) and Progenesis QI (Nonlinear Dynamics, UK). For each method, metabolite identities were confirmed using authentic reference standards.

#### Mouse retinal lipidomics

Metabolic profiling was performed by LC-MS on samples pooled from both retinas of P14 mice; samples were analyzed for fatty acylcarnitine species and beta-oxidation intermediate metabolites. Retinas were harvested after sacrificing P14 mice and immediately snap-frozen in liquid nitrogen. The acylcarnitines were extracted by protein precipitation (PP) and derivatized with butanol-HCl for analysis by LC/MS/MS. The samples were diluted with 400 µl solution of methanol with eight ISTD. After 1.5 minutes of homogenization in a bead Ruptor12 from Omni International Inc., samples were centrifuged for 5 minutes at 12000 rpm. The supernatant was transferred in a 2 ml polypropylene tube for evaporation under nitrogen. When dry, 100 µl of butanol-HCl (3N) is used for butanolysis performed by heating at 55°C for 20 min. After evaporation under nitrogen, the residue was reconstituted in 100 µl of mobile phase (ACN:H2O 80:20 with 0.05% of formic acid). Twenty microliters were injected in flow injection onto an Alliance 2795 LC coupled to a Quattro micro MS/MS (Waters). Data were recorded in positive electrospray ionization and analyzed with Neolynx from Waters Corp.

#### Metabolomics data analysis

Human metabolomics profiles were analyzed using MetaboAnalyst^3^. Data were filtered based on the interquartile range for further statistical comparisons. Outliers detected by the Random Forest algorithm were removed, and PCA, t-test, AUC test, and Metabolite Set Enrichment Analysis were performed on auto-scaled data previously normalized on internal standard (Phe-d8). Mouse acylcarnitine profiling was performed on unfiltered, auto-scaled values from MetaboAnalyst^3^ followed by PCA and hierarchical clustering (dendrogram and heatmap).

#### Droplet Sequencing (Drop-seq)

Drop-seq procedure was done on endothelial enriched cell suspension isolated from mouse retina. Briefly, single-cell suspensions were prepared from P14 and P17 retina, as reported^4^, through successive steps of digestion (using papain solution; Worthington, LK003150), trituration, and filtration to obtain a final concentration of 120 cells/μl. According to the manufacturer’s protocol, endothelial cells were enriched using CD31 microbeads (Miltenyi Biotech, 130-097-418). Droplet generation and cDNA libraries were performed as described in the Drop-seq procedure^4^, and sequencing was done on Illumina NextSeq 500. Unique molecular identifier (UMI) counts for WT and/or *Sirt3^−/−^* scRNAseq replicates were merged into one single Digital Gene Expression (DGE) matrix and processed using the “Seurat” package^5^. Cells expressing less than 100 genes and more than 10% of mitochondrial genes were filtered out. Single-cell transcriptomes were normalized by dividing by the total number of UMIs per cell and then multiplying by 10,000. All calculations and data were then performed in log space. PCA analysis on the most variable genes in the DGE matrix identified 20 significant PCs, which served as input for Uniform Manifold Approximation and Projection (UMAP). We used a density clustering approach to identify putative cell types on the embedded map and computed average gene expression for each identified cluster based on Euclidean distances. We then compared each cluster to identify marker genes differentially expressed across clusters. Cell type-specific transcriptomic differences between conditions, time points, and genotypes were statistically compared using a negative binomial model. Seurat visualization tools included Ridge Plot, Dot Plot, and UMAP Plot. Single-cell gene expression profiles from each separate cell type identified by scRNAseq were further analyzed using ggplot2^6^, Gene Set Variation Analysis^7^, EnrichR^8^, AUGUR^9^, Psupertime^10^, NicheNet^11^, SCENIC^12^ and Qusage^13^.

#### RNAscope *in situ* hybridization and immunohistochemistry

Experiments were performed per manufacturer instructions on fixed retinal cryosections (12 μm) using the RNAscope 2.5 chromogenic duplex assay (ACD bio-techne). WT and *Sirt3^−/−^* eyes at P17 were fixed in PFA 4% (24 h) at 4°C. Eyes were then immersed in a sucrose gradient (10 to 30%), frozen in optimal cutting temperature compound (Surgipath FSC22 Clear, Leica), and stored at -80 °C until use. *Aqp1* and *Hadha* mRNA expression were targeted using probes designed by ACD Bio-techne. For immunofluorescence, WT and *Sirt3^−/−^* eyes at P17 were fixed in PFA 4%, then the retina was dissected and stained overnight in blocking solution (4°C) with primary antibodies against ESM1 (1:100, R&D Systems, AF1999) or AQP1 (1:100, Abcam, ab125041), followed by secondary antibodies in blocking solution for 2 hours RT before flat mounting. RNAscope and IF retina were counterstained with isolectin B4 (Alexa Fluor 594, I21413; Molecular Probes). Retinas were then imaged with a Leica DME-6 microscope using a 20X dry objective (HC PL APO 20X/0.75 CS2) and analyzed with ImageJ (NIH).

#### Quantification of neovascularization (NV) and vaso-obliterated (VO) areas

Eyes were fixed for 1 hour in 4 % paraformaldehyde at room temperature (RT). Retinas were dissected and stained overnight at RT with fluoresceinated isolectin B4 (Alexa Fluor 594, I21413; Molecular Probes) in 1 mM CaCl_2_ in PBS. Lectin-stained retinas were whole-mounted onto Superfrost Plus microscope slides (Fisher Scientific) with the photoreceptor side down and embedded in SlowFade Antifade Reagent (Invitrogen). Stained retinas were flat-mounted, and the pictures were taken with an epifluorescence slide scanner, Zeiss Axio Scan.Z1 (Zeiss) at 10x magnification. The neovascular (NV) and vaso-obliterated (VO) areas were quantified using the SWIFT_NV program^14^.

#### Extracellular acidification rate

Extracellular acidification rate (ECAR) was measured using a Seahorse Analyzer XFe 96. Whole retinas were isolated and dissected in DMEM d5030 medium (supplemented with 2 mM glucose, 2 mM glutamine, and 1 mM sodium pyruvate), and retinas were flat mounted on nitrocellulose membrane. Four 1.5 mm retinal punches were collected per retina and were loaded onto 96 well spheroid plates. Retinal punches (adhered to nitrocellulose membrane) were incubated in assay medium (DMEM d5030, 12 mM Glucose, 2 mM Glutamine, 1 mM Sodium Pyruvate, 1 mM Hepes, 0.5mM carnitine, pH 7.4). Basal ECAR measurements were collected over the initial 30 minutes of the experiments (5 time points) because of the limited viability of retinal explants.

#### Retinal glucose uptake assay

Consumption of glucose was measured using 2-[^3^H]deoxyglucose (DG) as we reported^15^. Mice received a daily intraperitoneal injection of 0.5 µL of 2-[^3^H]DG (1 µCi / µ L) from P12 to P17. Animals were sacrificed at P17, 60 minutes after the last injection, to insure cellular incorporation of the radiotracer. Retinas were collected and homogenized in a scintillation cocktail (Ecolite +; MP Biomedicals), and beta counts were measured using an LS6500 Multipurpose Scintillation Counter (Beckman). Disintegrations per minute (DPM) counts were normalized to the total injected dose.

### QUANTIFICATION AND STATISTICAL ANALYSIS

Statistical analysis was performed using GraphPad Prism (10.0.3) for Mac (GraphPad Software) or R statistical programming language (4.0.3, R-project.org). Exact n, statistical test, and significance are reported in the Figure Legends. Data are presented as means ± the standard error of the mean (s.e.m.). Between subject statistical tests were completed with two-tailed independent Student’s *t*-tests. Comparisons between groups were made using 1-way or 2-way ANOVA followed by the post hoc Tukey or Bonferroni multiple comparisons test among means. The D’Agostino-Pearson or Kolmogorov-Smirnov normality tests were used to confirm a normal distribution. Animals were not randomized but quantification of the data was performed in a blinded fashion when possible. Outlier results were identified and removed using the ROUT method (Q1%) in GraphPad Prism. Results are presented as mean ± SEM. * P < 0.05, ** P < 0.01, *** P < 0.001, **** P < 0.0001. *P* < 0.05 was considered statistically significant.

## SUPPLEMENTAL TABLE LEGEND

**Supplemental Table 1.**
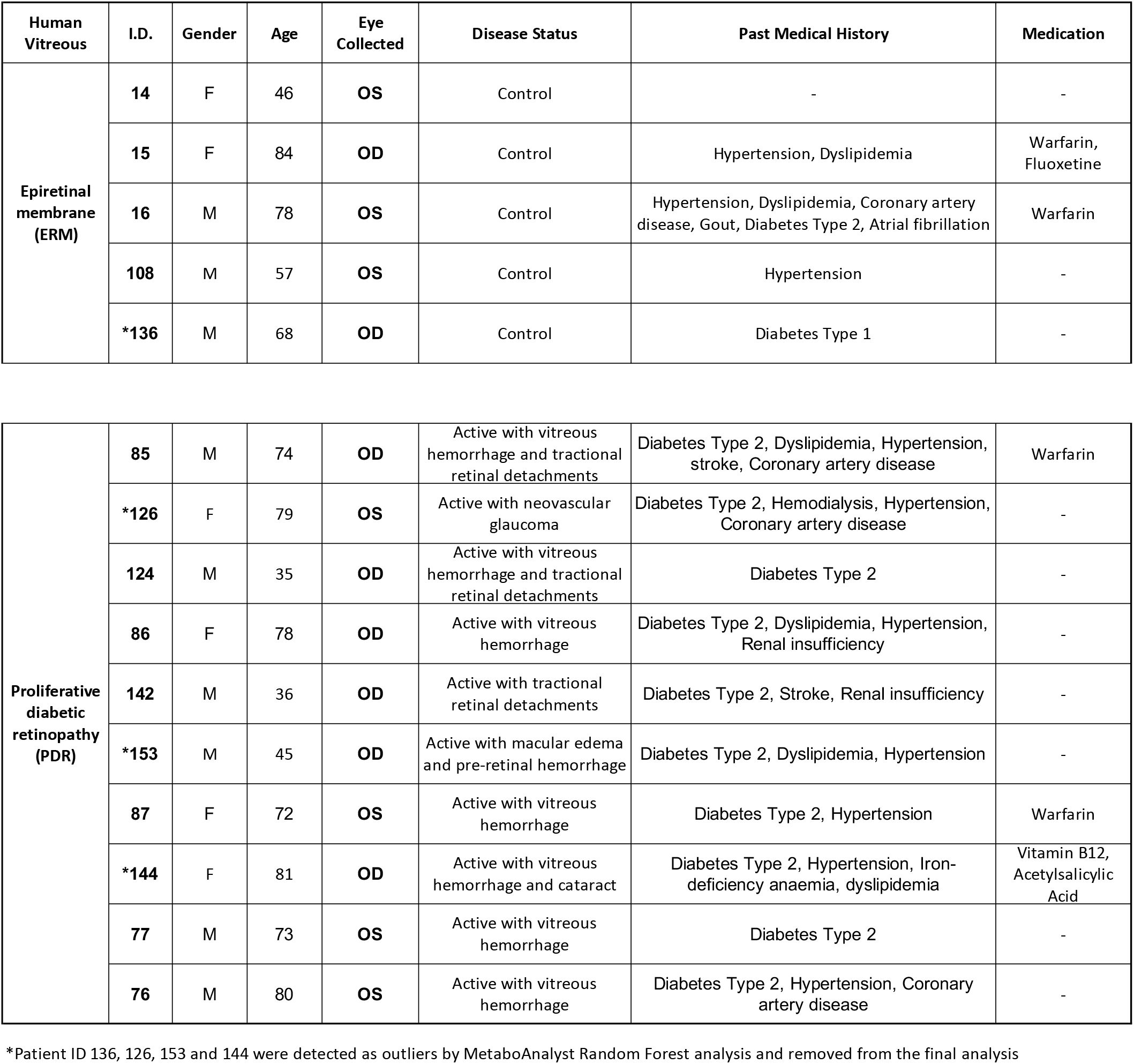
Clinical information of human subjects. Vitreous was collected either in the right (OD) or left (OS) eye of patients with epiretinal membrane (control) and proliferative diabetic retinopathy (PDR).

| Human Vitreous | I.D. | Gender | Age | Eye Collected | Disease Status | Past Medical History | Medication |
| --- | --- | --- | --- | --- | --- | --- | --- |
| Epiretinal membrane (ERM) | 14 | F | 46 | OS | Control | - | - |
|  | 15 | F | 84 | OD | Control | Hypertension, Dyslipidemia | Warfarin, Fluoxetine |
|  | 16 | M | 78 | OS | Control | Hypertension, Dyslipidemia, Coronary artery disease, Gout, Diabetes Type 2, Atrial fibrillation | Warfarin |
|  | 108 | M | 57 | OS | Control | Hypertension | - |
|  | *136 | M | 68 | OD | Control | Diabetes Type 1 | - |
| Proliferative diabetic retinopathy (PDR) | 85 | M | 74 | OD | Active with vitreous hemorrhage and tractional retinal detachments | Diabetes Type 2, Dyslipidemia, Hypertension, stroke, Coronary artery disease | Warfarin |
|  | *126 | F | 79 | OS | Active with neovascular glaucoma | Diabetes Type 2, Hemodialysis, Hypertension, Coronary artery disease | - |
|  | 124 | M | 35 | OD | Active with vitreous hemorrhage and tractional retinal detachments | Diabetes Type 2 | - |
|  | 86 | F | 78 | OD | Active with vitreous hemorrhage | Diabetes Type 2, Dyslipidemia, Hypertension, Renal insufficiency | - |
|  | 142 | M | 36 | OD | Active with tractional retinal detachments | Diabetes Type 2, Stroke, Renal insufficiency | - |
|  | *153 | M | 45 | OD | Active with macular edema and pre-retinal hemorrhage | Diabetes Type 2, Dyslipidemia, Hypertension | - |
|  | 87 | F | 72 | OS | Active with vitreous hemorrhage | Diabetes Type 2, Hypertension | Warfarin |
|  | *144 | F | 81 | OD | Active with vitreous hemorrhage and cataract | Diabetes Type 2, Hypertension, Iron-deficiency anaemia, dyslipidemia | Vitamin B12, Acetylsalicylic Acid |
|  | 77 | M | 73 | OS | Active with vitreous hemorrhage | Diabetes Type 2 | - |
|  | 76 | M | 80 | OS | Active with vitreous hemorrhage | Diabetes Type 2, Hypertension, Coronary artery disease | - |

\*Patient ID 136, 126, 153 and 144 were detected as outliers by MetaboAnalyst Random Forest analysis and removed from the final analysis

## SUPPLEMENTAL FIGURE LEGENDS

**Supplemental Figure 1.**
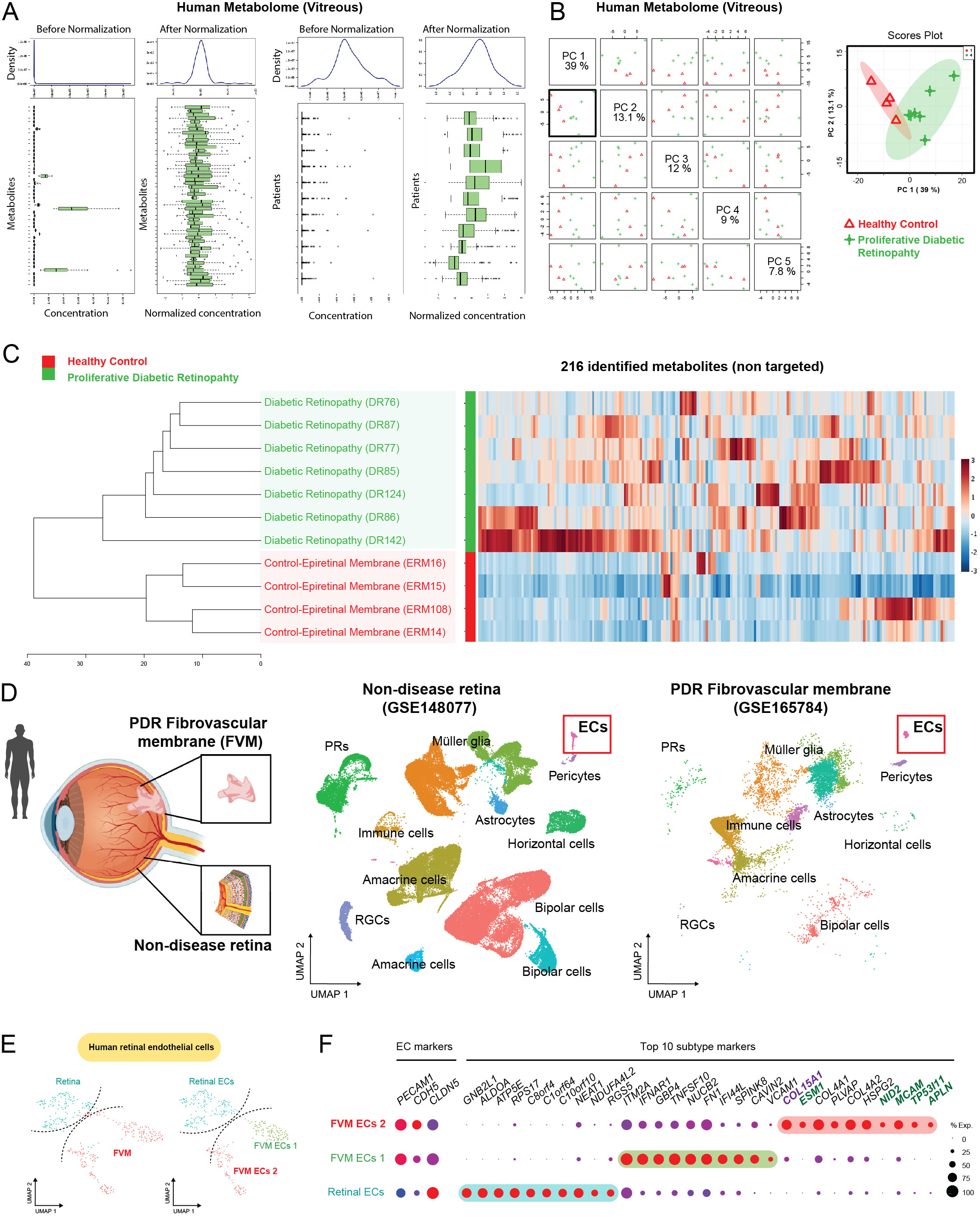
**A.** Normalization of vitreous metabolomics data (see method) showing the distribution of metabolites (left) and patient samples (right). **B.** Principal component analysis of vitreous metabolomics data across control (red, n=4) and proliferative diabetic retinopathy (green, n=7) patients. **C.** Heatmap clustering of all metabolites across control (red) and proliferative diabetic retinopathy (green) patients. **D.** UMAP of publicly obtained scRNAseq integrated datasets of human post-mortem retina (GSE148007) and fibrovascular membranes (FVM) from human patients with proliferative diabetic retinopathy (GSE165784). **E.** UMAP of endothelial subclusters from control and FVM derived-human retinal sample (EC subset from S1D). **F.** Dotplot of top 10 markers for the 3 identified EC subclusters, showing the presence of an FVM endothelial subcluster (FVM ECs 2) expressing gene markers of neovascular ECs (ESM1, TP53I11, APLN, PLVAP).

**Supplemental Figure 2.**
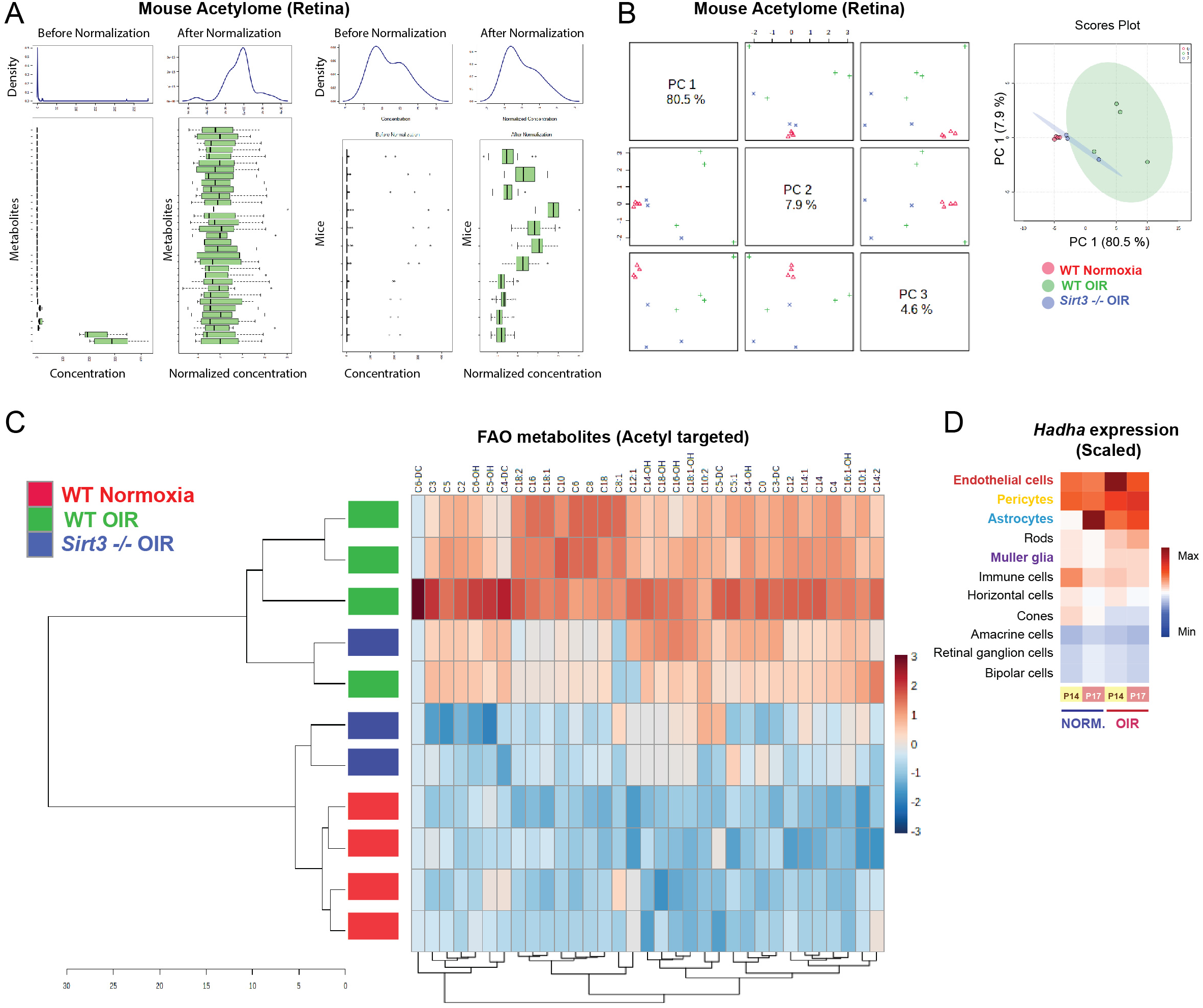
**A.** Normalization of retinal lipidomics data showing the distribution of metabolites (left) and retinal samples (right) from WT normoxia (n=4), WT OIR (n=4), and *Sirt3^-/-^* OIR (n=3) P14 mice. **B.** Principal component analysis of retinal lipidomics data across WT normoxia (red), WT OIR (green), and *Sirt3^-/-^* OIR (blue) P14 mice. **C.** Heatmap clustering of retinal samples based on all lipidomics metabolites for WT normoxia (red), WT OIR (green), and *Sirt3^-/-^* OIR (blue) P14 mice. **D.** Expression levels of Hadha (Hydroxyacyl-CoA Dehydrogenase Trifunctional Multienzyme Complex Subunit Alpha), which encodes a subunit of the mitochondrial trifunctional protein essential for acyl-carnitine b-oxidation, from single-cell RNAseq analysis of normoxic and OIR mouse retina at P14 and P17 (GEO accession number GSE150703).

**Supplemental Figure 3.**
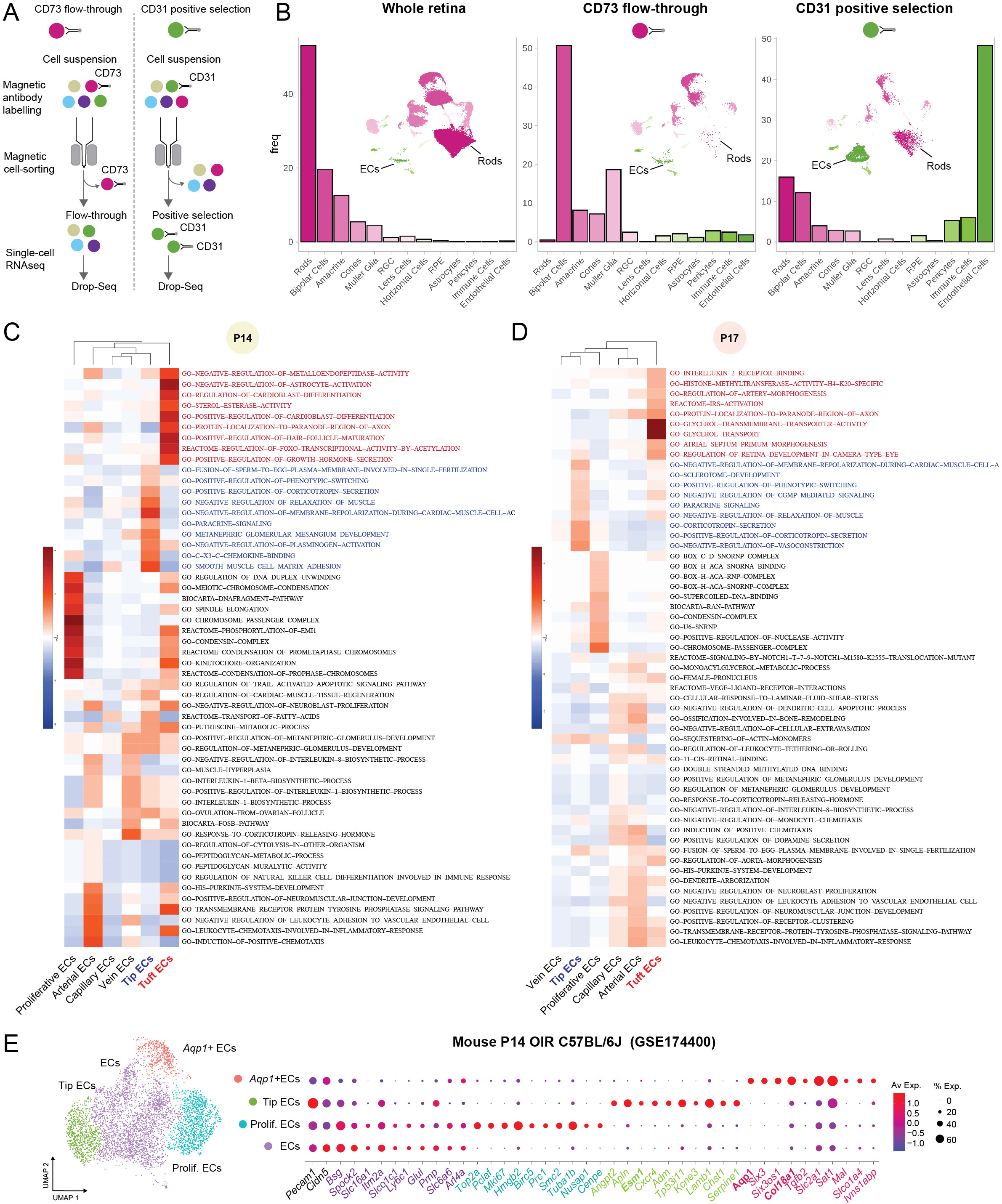
**A.** Two different approaches to enrich cell types of interest for single-cell RNAseq analysis, using magnetic cell isolation technology (MACS, Miltenyi Biotec). Retinal cell suspension can either be depleted of rods (CD73 antibody) or enriched for endothelial cells (CD31 antibody) **B.** Single-cell RNAseq illustrating the cell type distribution using the whole retina compared to CD73 depleted or CD31 enriched cell suspensions. **C and D.** Heatmap of pathway enrichment based on GSVA analysis of scRNAseq data showing the normalized GSVA score for the most enriched pathways in each endothelial subtype of OIR retina at P14 and P17. **E.** UMAP of publicly obtained scRNAseq of P14 retina from C57B6/6J mice exposed to OIR (GSE174400) showing the presence of tuft EC subcluster (Aqp1, Col18a1).

**Supplemental Figure 4.**
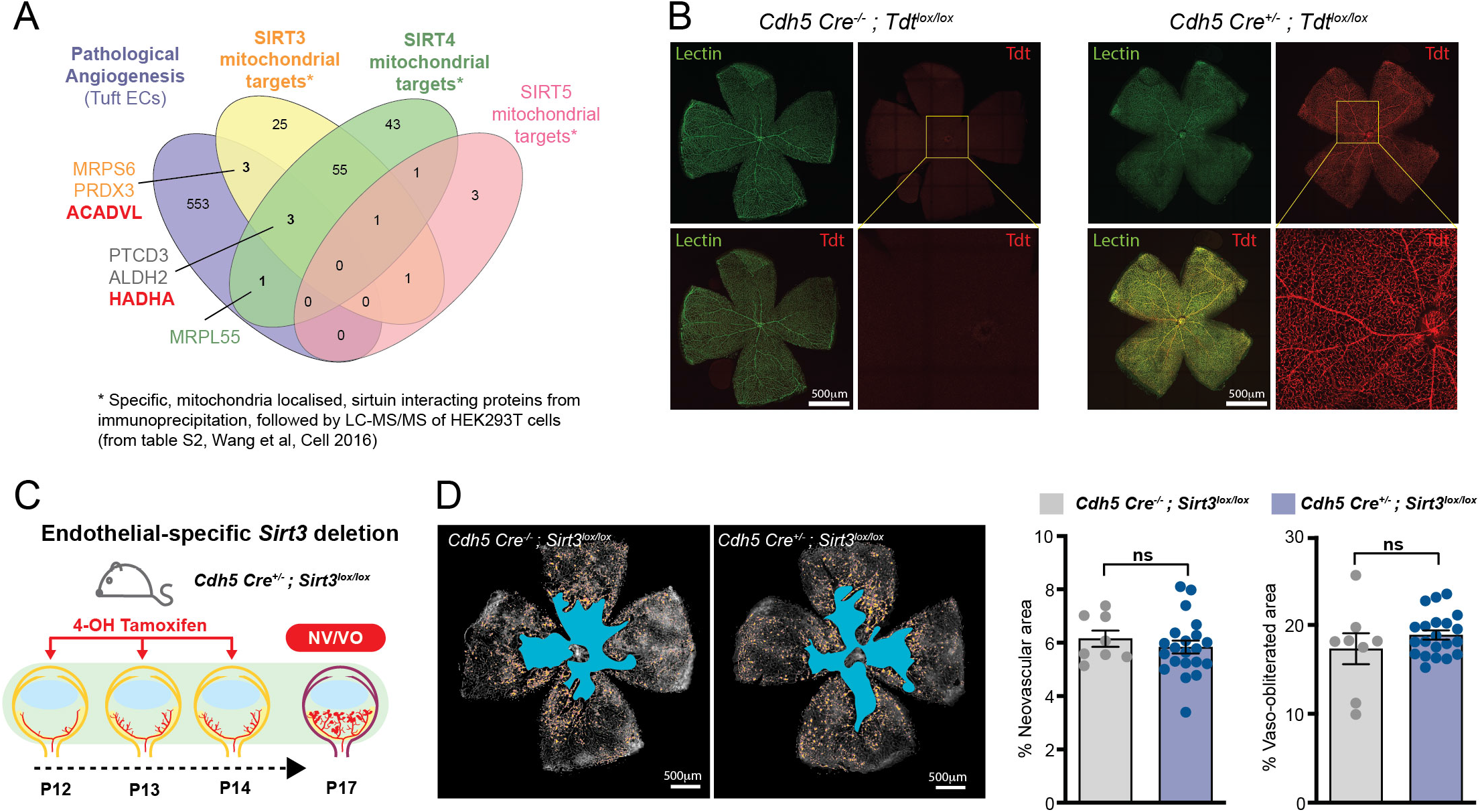
**A.** Overlap between mitochondrial sirtuins (SIRT 3,4,5) interacting proteins (from HEK293T cells) and differentially expressed genes (DEG) in tuft and tip EC from OIR retinas (P17); 6 DEG in tuft EC, including ACADVL and HADHA, were proteins known to directly interact with SIRT3. **B.** Retinal flatmounts of tamoxifen-injected Cdh5-CreERT2 expressing mice, or not, and bred with tdTomato*^lox/lox^* (Tdt) reporter mice. Endothelial expression of Cdh5-CreERT2 by 4-OH tamoxifen (red) localized to retinal vessels (lectin, green). **C.** Graphical representation of the experimental strategy to induce EC-specific deletion of *Sirt3* during the neovascularization period of OIR. Cdh5-CreERT2^-/-^ ; Sirt3^lox/lox^ and Cdh5-CreERT2^+/-^ ; Sirt3^lox/lox^ mice were treated from P12 to P14 with daily injections of 4-OH Tamoxifen (5 μg). **D.** Lectin-stained retinal flat-mount of P17 Cdh5-CreERT2^-/-^ ; Sirt3^lox/lox^ (n=8 retinas) and Cdh5-CreERT2^+/-^ ; Sirt3^lox/lox^ mice (n=21 retinas) exposed to the OIR model; vaso-obliterated (VO, blue) and neovascular (NV, yellow) areas are highlighted. Bar graph represents VO and NV areas relative to total retinal areas (unpaired Student t-test, ns: P-value > 0.05).

**Supplemental Figure 5.**
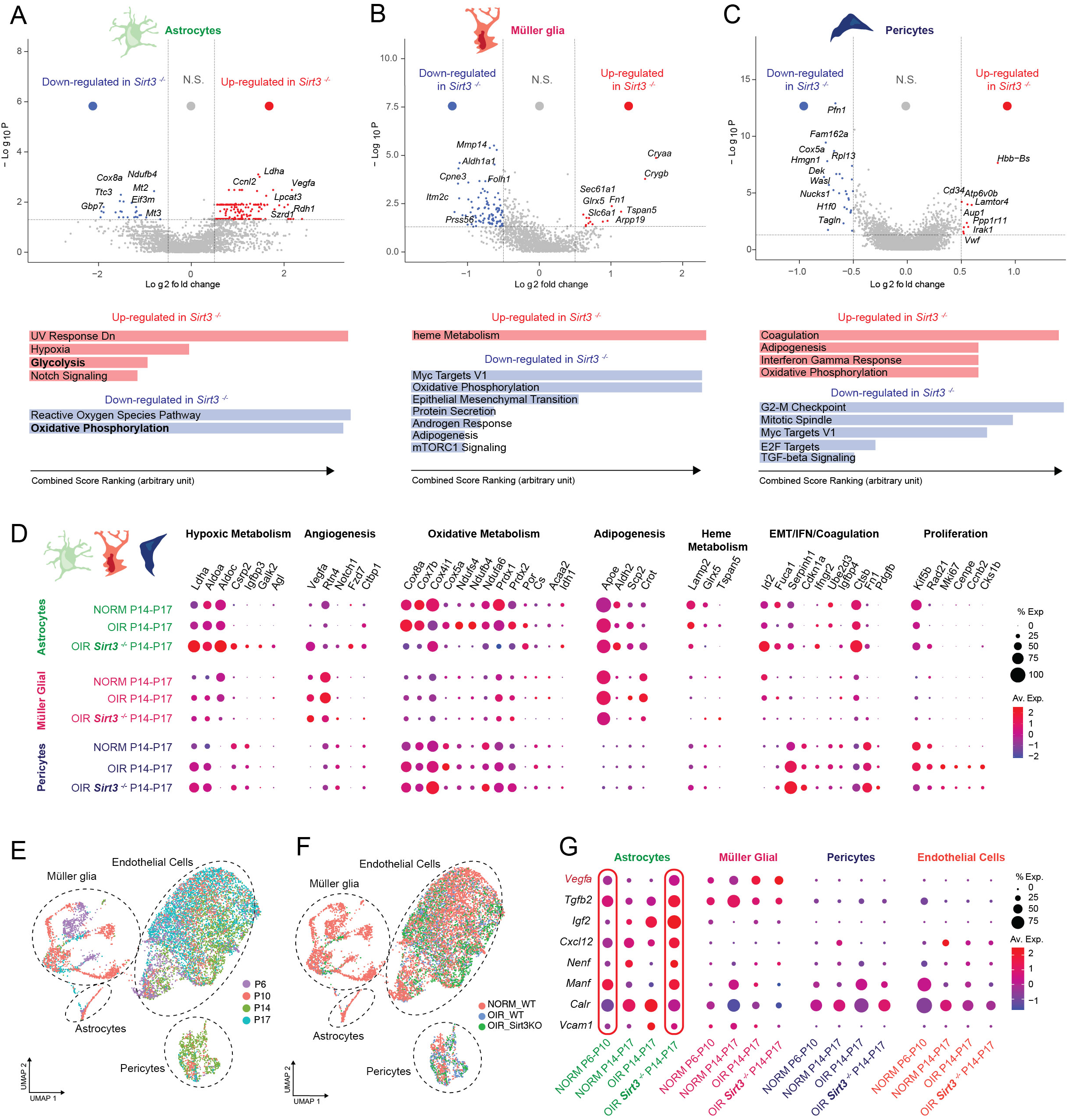
**A-C.** Volcano plot of differentially expressed genes between OIR WT and *Sirt3^-/-^* astrocytes, Müller glia, and pericytes (P value <0.05, absolute log2FC > 0.25, non-parametric Wilcoxon rank sum test, excluding genes expressed in less than 10% of cells in the given cell type), and their associated pathways from EnrichR. **D.** Dotplot of differentially expressed genes between WT and *Sirt3^-/-^* in cells of the neurovascular unit in OIR compared to normoxia. **E and F.** UMAP of integrated single-cell RNAseq from publically available mouse retinal P6-10 dataset (10X, GSE175895) with the current P14-17 dataset. **G.** Dot plot representing the expression of selected ligands that were differentially expressed between cells of the vascular unit in WT and Sirt3-/-retinas using NicheNet analysis during development (P6-10) and compared to OIR (P14-17).

**Supplemental Figure 6.**
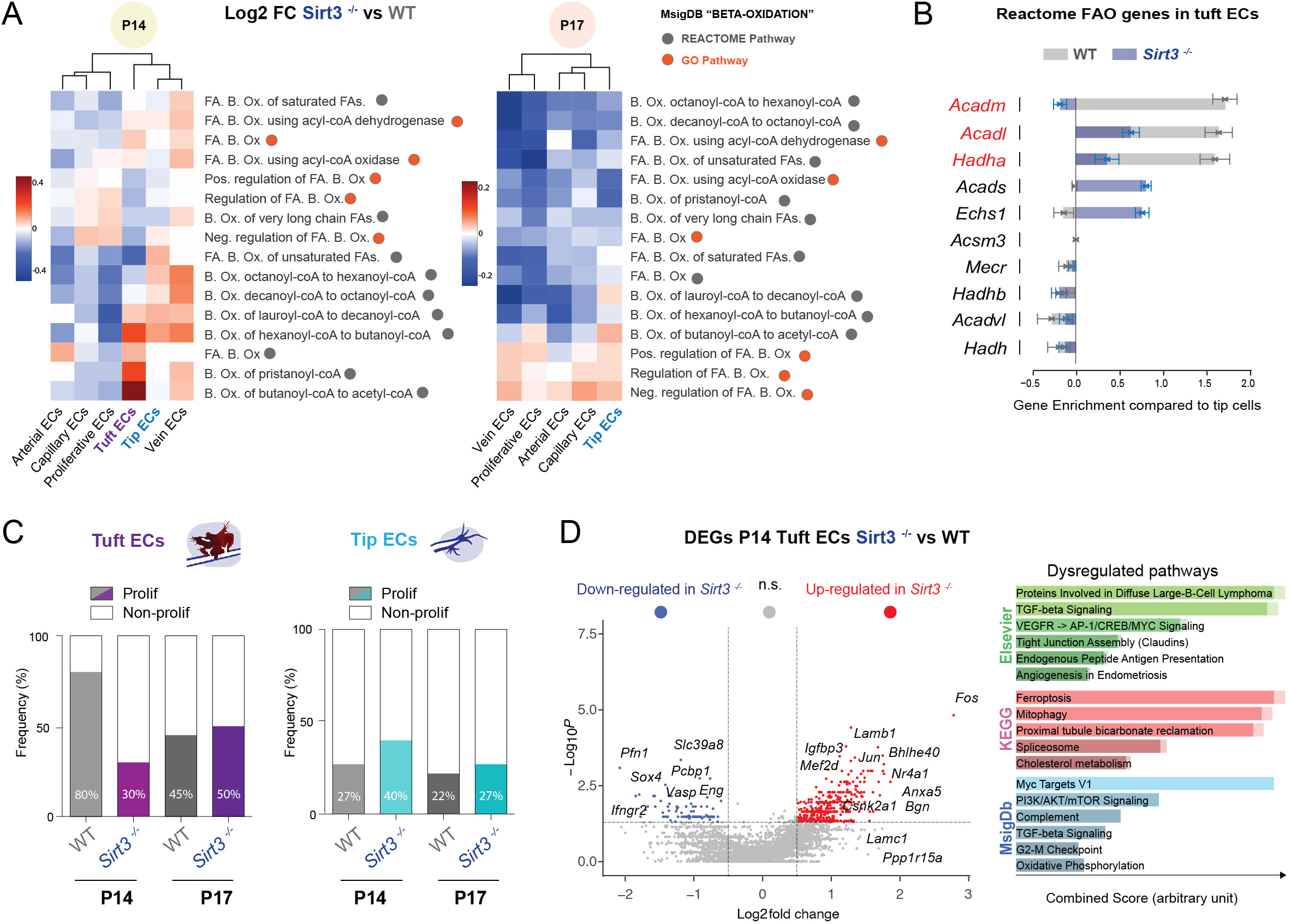
**A.** Heatmap of normalized GSVA score log2 fold-change (FC) between OIR WT and *Sirt3^-/-^* EC subtypes for several FAO pathways from the GO and REACTOME Molecular Signature database at P14 and P17. **B.** Comparison by Qusage of pathway enrichment in tuft EC relative to tip cells for the REACTOME Fatty-acid oxidation pathways in P14 OIR WT and *Sirt3^-/-^* retina. Qusage Gene Set Enrichment-type test uses the Variance Inflation Factor technique and quantification of gene set activity with a complete probability density function. **C.** Percent of non-proliferative (G1) and proliferative cells (S, G2/M) amongst tip and tuft EC per genotype and time point. **D.** Volcano plot of differentially expressed genes between P14 OIR WT and *Sirt3^-/-^* tuft EC (P value <0.05, absolute log2FC > 0.5, non-parametric Wilcoxon rank sum test, excluding genes expressed in less than 10% of cells in the given cell type), with corresponding enriched pathways from EnrichR.

